# Supramolecular cylinders target bulge structures in the 5’ UTR of the RNA genome of SARS-CoV-2 and inhibit viral replication

**DOI:** 10.1101/2021.03.30.437757

**Authors:** Lazaros Melidis, Harriet J. Hill, Nicholas J. Coltman, Scott P. Davies, Kinga Winczura, Tasha Chauhan, James S. Craig, Aditya Garai, Catherine A..J. Hooper, Ross T. Egan, Jane A. McKeating, Nikolas J. Hodges, Zania Stamataki, Pawel Grzechnik, Michael J. Hannon

## Abstract

The untranslated regions (UTRs) of viral genomes contain a variety of conserved yet dynamic structures crucial for viral replication, providing drug targets for the development of broad spectrum anti-virals. We combine in vitro RNA analysis with Molecular Dynamics simulations to build the first 3D models of the structure and dynamics of key regions of the 5’ UTR of the SARS-CoV-2 genome. Furthermore, we determine the binding of metallo-supramolecular helicates (cylinders) to this RNA structure. These nano-size agents are uniquely able to thread through RNA junctions and we identify their binding to a 3-base bulge and the central cross 4-way junction located in the stem loop 5. Finally, we show these RNA-binding cylinders suppress SARS-CoV-2 replication, highlighting their potential as novel antiviral agents.

## Introduction

SARS-CoV-2 is a novel coronavirus that causes COVID-19 and as of 1st March 2021 there have been 113,267,303 recorded cases and 2,520,550 deaths worldwide[1]. Emerging so soon after other major coronavirus outbreaks (SARS, MERS), this global pandemic has highlighted the need for greater preparedness to tackle newly emergent viruses that may spread with lethal consequences. Fundamental understanding of viral processes needs to be coupled to the development of a variety of broad-acting antiviral strategies to interfere with these processes, in order to maximise the armory of drugs that we have available to treat novel pathogens. To date, antiviral drug designs have largely targeted viral proteins[2,3] especially those with enzymic functions such as proteases and polymerases[4,5]. An alternative approach is to target viral nucleic acid structures that are essential for replication. With current advances in sequencing technology, the sequence of a new virus can be identified within the first weeks of an outbreak, identifying both the protein coding regions and the untranslated regions (UTRs). The role of the UTRs is not completely understood for many viral families their conserved structures underline their functional importance. Where UTRs have been studied to determine function (retrovirus HIV-1[6,7], flavivirus[8-11], to a lesser extent coronavirus[12-14]) they have been shown to have dynamic structures essential for the viral replication[15,16].

These non-coding RNA regions are highly structured with multiple stem loops, bulges, crosses and pseudo-knots, with common structural elements seen in many viral UTRs. These structures play a role in RNA-RNA interactions (both within the viral genome and with host machinery) and in protein binding for the initiation of mRNA production, translation and viral replication. Moreover, these RNA structures may act as trans acting elements or mediate translational frameshifting, a common feature in viruses with plus-strand RNA genomes.

Nucleic acid sensors mediate the early detection and host response to virus infections, and recognise either viral nucleic acids or ‘unusual’ cellular nucleic acids present upon infection[17]. Sensors from the RIG-I-Like Receptor (RLR) family are key pattern recognition receptors for coronaviruses[18,19] which detect RNAs with specific structures such as 5’-triphosphate or 5’-diphosphate ends[20,21]. Therefore UTR structures within double-stranded viral RNA provide attractive drug targets, both for direct inhibition of viral replication[13] and induction of host innate immune responses.

Compared to protein- and DNA-recognition, RNA-recognition by drugs has been much less explored. Nucleic acid recognition often focuses on sequence recognition but for RNA, which folds into complex shapes, its structure provides an opportunity for specific targeting; indeed, it is the structure of the UTR that is conserved for function, rather than sequence. Small molecule libraries have been screened for RNA binding (analogous to protein drug screens)[22-24] and agents targeting RNA structures include small molecules that hydrogen bond within the heart of trinucleotide DNA/RNA repeats[25], and planar RNA quadruplex binders[26-31].

We have explored nano-size metallo-supramolecular cylinders (Fig.1A) as RNA-binding agents[32]. They are larger than traditional small molecules, with extensive aromatic surfaces to stack with the RNA bases (Fig. 1B) and cationic charge (4+) that ensure strong binding and excellent shape-fit for RNA cavities. We have characterized the binding of cylinders in an RNA 3-way junction[32] by crystallography (Fig. 1C) and showed analogous binding in an RNA bulge structure[33,34]. Furthermore, we demonstrated cylinder binding to an RNA 3-base bulge in the TAR region of the HIV-1 genome (located in its UTR), that prevented HIV-1 replication[34].

**Figure 1.**
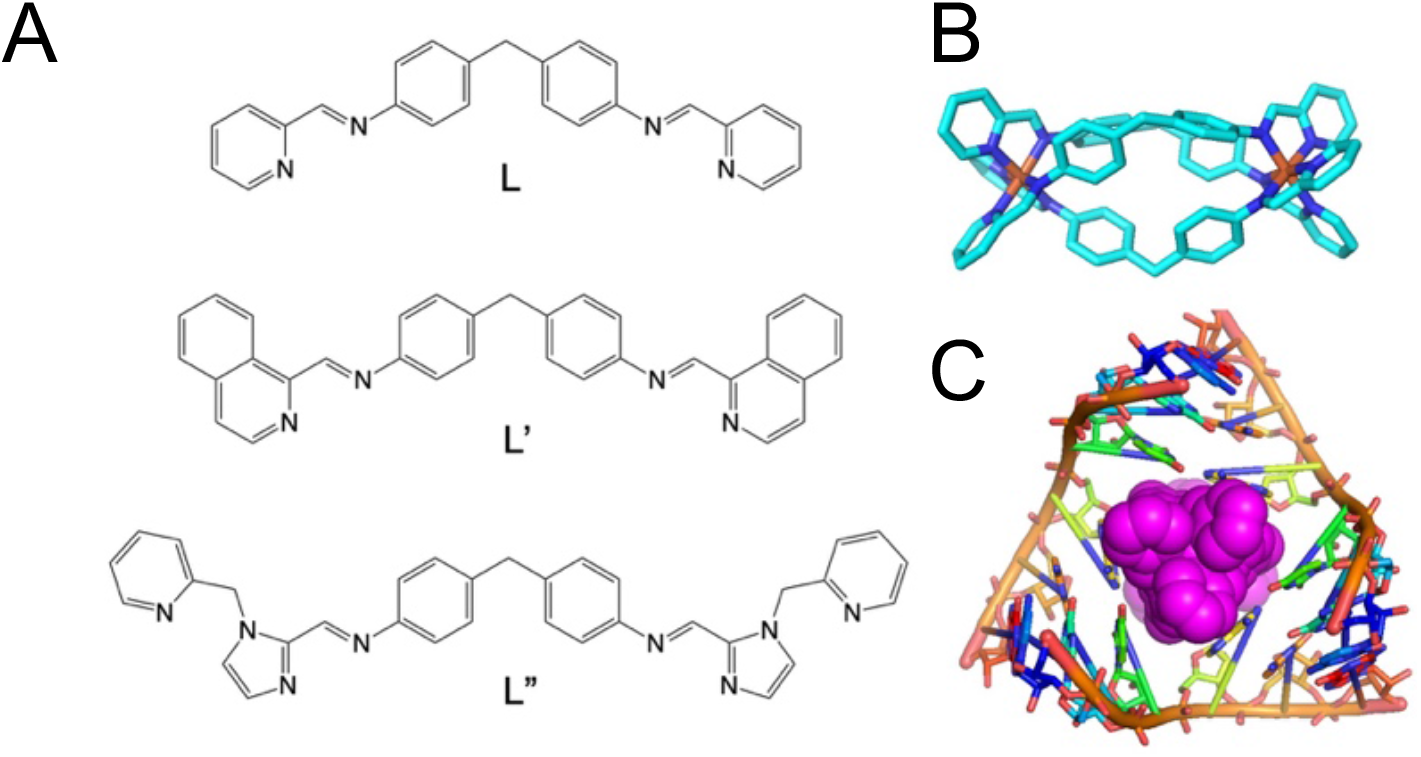
(A) Structure of the ligands used in this study. (B) Structure of the [Ni_2_L_3_]^4+^ cylinder of ligand L,^[35]^ L’ ^[36]^ and L’’ ^[37]^ form analogous cylinders that bear further aryl rings on their external surfaces. (C) View of the crystal structure of a cylinder bound in an RNA 3-way junction cavity from pdb 4JIY^[32]^ showing its unique binding.

Given this antiviral activity against HIV-1 we were interested to assess whether these cylinders would bind structures in the 5’ UTR of SARS-CoV-2. We now report combined modelling and biophysical approaches to define the 3D structures of the SARS-CoV-2 5’ UTR, and demonstrate cylinder binding to specific bulge structures in the 5’ UTR. Furthermore, we show that cylinders inhibit SARS-CoV-2 viral replication in cells.

## Results and discussion

To create a 3D dynamic model of the 5’ UTR from the published genome sequence[38] (original Wuhan strain, NC_045512), our approach was to predict the secondary structures in silico, obtain experimental evidence to verify these structures, and then model the tertiary structure and its dynamic behavior, again with experimental validation. RNA secondary structure prediction has improved dramatically over the last decade, with free energy approximations and machine learning algorithms available (adding to the attraction of the RNA as a rapid-response drug target). However, there are significant challenges with longer RNA sequences that can yield multiple distinct structures that occupy a small space in the energy landscape. We compared ∼10 folding prediction algorithms (see Supplementary Information) with many failing to cope well with the large size of the SARS-CoV-2 5’ UTR. Three representative predictions are shown in Figure 2. The free energy RNAfold[39] and Mxfold2[40] algorithms gave similar predictions, both akin to the known UTR structures of related coronaviruses[16][41], while the machine learning based VFold[42] gave a quite distinct structure.

**Figure 2.**
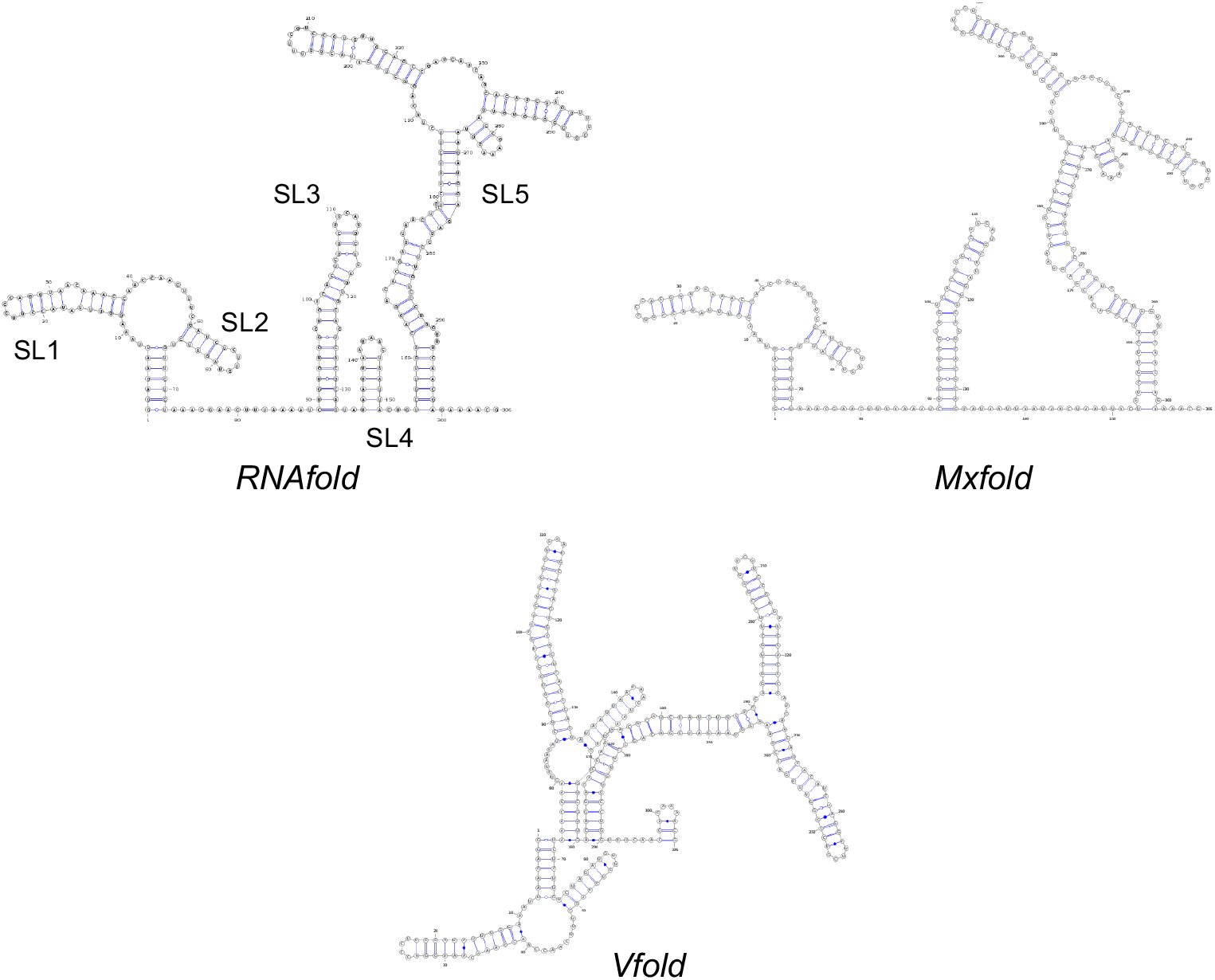
Secondary structure predictions of the UTR of SARS-CoV-2 using three different algorithms.

To experimentally probe the UTR, we used SHAPE, (Selective 2’-Hydroxyl Acylation Analyzed by Primer Extension Sequencing) analysis where the 5’ UTR RNA sequence was first folded in vitro and the open strand (non-duplex) RNA sites (e.g. single stranded, bulges, hairpins) acylated with 1-methyl-7-nitroisatoic anhydride (1M7). These sites were then identified through a reverse transcription reaction that generates DNA fragments which end at the 1M7 tagged sites and were analysed by gel electrophoresis (Fig 3A). Two primers (RT1 and RT2) conjugated with fluorescent IRDye700 were used to cover the whole 5’ UTR sequence. RT1 mapped the UTR from position +1 to +140, and RT2 the distal region of the UTR (+141 to +300). The results (summarized as a diagram in Fig. 3B) demonstrate that the RNAfold/Mxfold predicted structures best represent that formed in vitro. In particular, the long run of acylation around position G confirms that the Vfold prediction does not adequately describe the experimental data. The additional stem-loop (SL4) predicted by RNAfold but not Mxfold is acylated (region K) which suggests that if such a stem loop forms it may be transient. Recent studies of the whole RNA viral genome *in cellulo* by Miska[43] (COMRADES assay) and Pyle[44,45] (long amplicons with SHAPE-MaP) show dynamic folding and interaction between the 5’ UTR and 3’ UTR, but that these key stem-loop structures (SL1,2,3,5; depicted in Fig2) are retained, affording further support and confidence that our in vitro findings are physiologically relevant.

**Figure 3.**
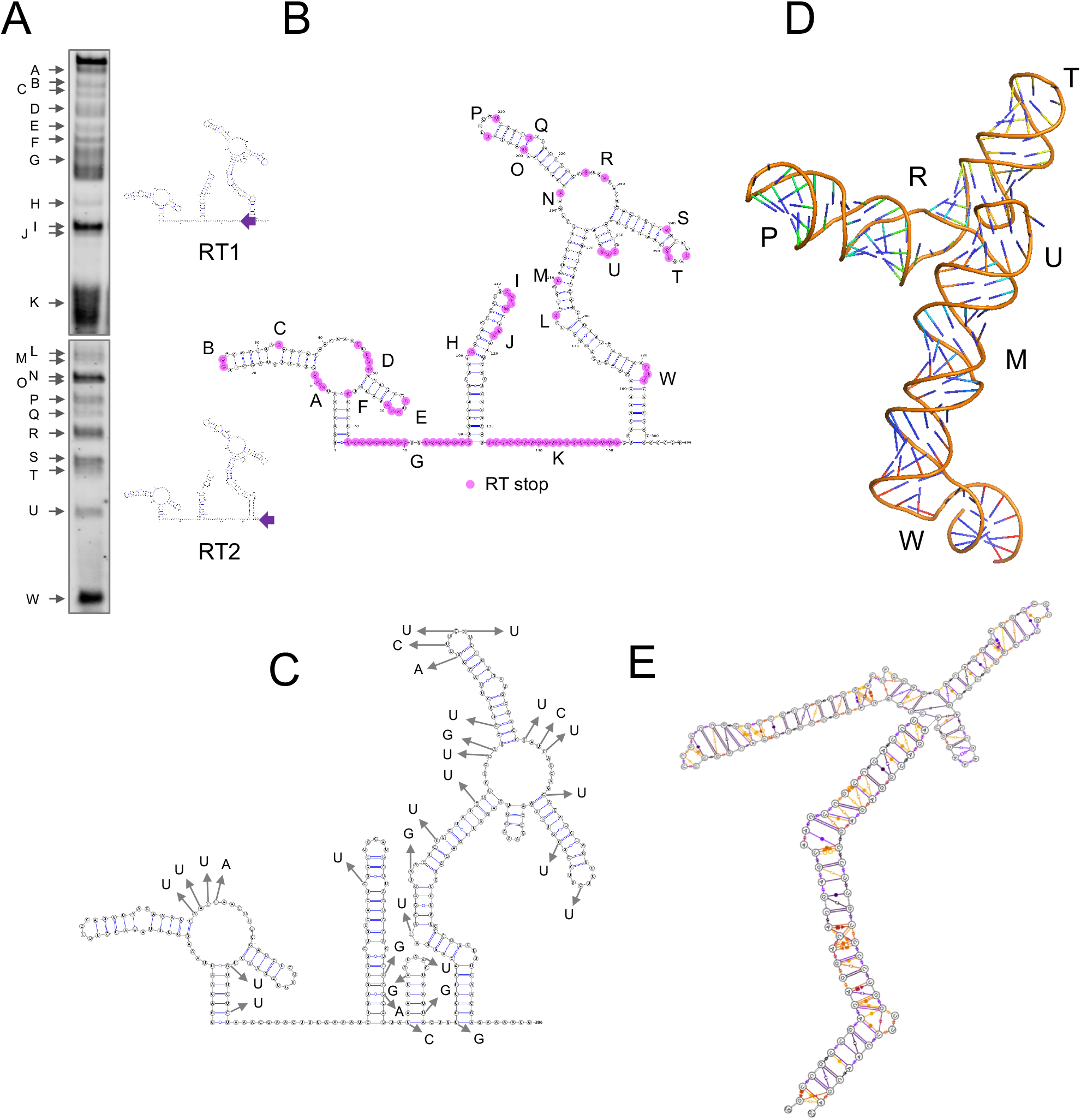
The structure of the SARS-CoV-2 5’ UTR. (A) RNA SHAPE gel results. Two IRD700 RT primers were used; RT2 primer maps the whole sequence however longer molecules are not very well separated by electrophoresis so additional RT1 was used to map the 5’ region in more detail (diagrams showing positions of the primers are included). (B) SARS-CoV-2 5’ UTR secondary structure showing the acylated nucleotides revealed by RT stops as purple dots. Open structures are labelled A-W. (C) Positions of SNPs observed in SARS-CoV-2 viral sequences to 7 Jan 21. See also Fig. S6 for overlay of 3B and 3C (D) Snapshot of the dynamic three-dimensional structure of the SL5 RNA from MD simulations (E) Leontis Westhoff diagrams highlighting the dynamic base-pairing within the structure.

The extensive whole-genome sequencing of SARS-CoV-2 affords the opportunity to monitor the single nucleotide polymorphism (SNPs) mutations in the 5’ UTR. We examined the available sequences in the gisaid[46] that were deposited before 7 January 2021 that contained a complete 5’ UTR. Interestingly the positions of SNPs within the UTR (Fig. 3C) often occur near the acylated positions in our SHAPE experiment (Fig. 3B, S6), suggesting that positions where the nucleotide has greater flexibility and hence less structural importance for the UTR are more likely to be substituted. Although not corrected for frequency, it is interesting to note that around 60% (19/31) of the SNP sites identified to date involve replacement with a U residue, with the largest subset (11/31) being a C-U mutation (Fig. S6). As anticipated the mutations will not affect the key structures of the 5’ UTR.

After identifying the distinct stems loops (SLn) that were conserved throughout the results from the secondary structure prediction, we attempted the more challenging step of creating a 3-dimensional representation of the structure. We focused on SL3 and SL5 as they have a variety of different structural features including bulges and loops. Although the exact structure/function of SL5 is not yet determined (to our knowledge), it contains the initiation codon and is similar to the SL5 of SARS-CoV-1[12,13], suggesting a functional role. Understanding the tertiary structure and behaviour from the sequence, is more complicated than predicting the secondary sequence since RNA is an inherently flexible molecule and a single static conformation will not be sufficient to understand the binding properties. Recent advances in molecular dynamics parameterization of RNA and wider availability of high-performance computer facilities can provide new insights into the dynamic structure of the RNA and show the key regions of flexibility - usually bulges and junctions, where both the secondary and tertiary structure is highly dynamic. After creating initial models using the short list of open-source software available, the ROSETTA platform (FARFAR2)[41,47] gave a starting structure most consistent with the SHAPE analysis (notably the SL5 junction point having nucleotide interactions rather than being very open), so we explored the dynamics around this central structure.

We employed the recent RNA-force field developed by Mathews[48,49], which retains NMR characteristics of RNA structures even in non-minimum starting conformations, and coupled it with Markov state modeling[50] to analyse the conformational space accessed across different simulations. We started with 3 independent 1 microsecond molecular dynamics simulations of the SL5 alone, and then performed additional 1microsecond simulations with both enantiomers of the cylinder (three runs of at least 1 microseconds each; with parent cylinder and both enantiomers) to identify RNA regions that can be recognised by the cylinder. The simulations total 9µs. Additionally, Markov state modelling revealed micro states where the cylinder can be positioned within the RNA helix in the bulge regions. We also performed simulations on the SL3, comprising overall 4µs.

Just as for the secondary structure predictions, the observations in the molecular dynamics of SL5 were verified experimentally by the SHAPE results, and by using these techniques in concert we gain a molecular level understanding of the three dimensional structure and dynamic behaviour of the RNA (Fig. 3.C, E) and how the cylinders bind.

Considering the SL5 RNA in absence of cylinder, molecular dynamics reveal the following features of the stem: (a) There is a bulge at G138-U140 which is highly flexible with a lot of transient stacking between its bases (region W in Fig. 3.B). G138 base pairing with C10 elongates the bulge forcing U139-U141 to point outwards of the helical axis. This is seen experimentally in SHAPE. This sharp twist of the backbone often creates a bend to the stem. (b) There is a mismatch at C15 (halfway between regions L and W) however there are many transient non-Watson-Crick base pairings between A14-A16 and C133 and those nucleotides did not produce a SHAPE signal; i.e. there is no significant bulge or base flipping outwards and the helix is contiguous (c) The next bulge (U21-U25; region L) is different. Relative stability is provided by three G:C base pairs (G20 :C128, C24:G126, C26:G124), causing flagging out of A23 as seen on SHAPE (region M). (d) At the 4-way junction (regions N, R) the base pairings (‘CUG’36-37 and ‘CAG’78-80) hold throughout the simulation (3µs) creating an additional 7 nucleotide bulge on SL5a (G72-A79) where on the opposite strand there are only C38 A39. Although C38 remains stacked to G37 and transiently binds nucleotides of the opposite strand A39 lacks both strong stacking or base pairing, therefore it can be seen on SHAPE. The junction is less open (i.e contains more pairing) than the secondary structure prediction and this is reflected in the SHAPE experiment where there is only limited acylation. (e) Higher up on the SL5a CG Watson-Crick (WC) pairs create rigidity which stops on the U47, which stacks strongly on C46 allowing stable non WC base paring with U67 but leaves U48 randomly pairing U66 and G66 (region O,Q). U48 and G66 are both identified by SHAPE. The stem closes with strong CG pairings and a short loop (region P), whose bending exposes U91 and U96 and they are identified by SHAPE. (f) On SL5b five CG pairs add rigidity allowing/stabilising non WC pairings. However, between C86:G100 and G89:C98 (region S) there is an additional base and as U87 and G99 strongly stack on the C86:G100 A88 is exposed and tagged by SHAPE. On the loop (region T) stacking continues strongly up to U92 and G95 creating a tight bend exposing U93. (g) The short SL5c is also stabilised by 2 CG pairs and all three A residues are stacked together but point outwards of the stem (region U).

These combined simulation/experimental pictures of the RNA dynamics were then complemented by analogous SHAPE experiments and MD simulations of the SL5 RNA in the presence of the [Fe_2_L_3_]^4+^ cylinder (Fig. 4). Four batches of simulations were carried out in the presence of cylinder; for each enantiomer of the cylinder and with the cylinders positioned either away from the RNA or inside the bulges. Importantly, the MD simulations locate the cylinder binding sites on SL5 at the same positions that are affected experimentally in the SHAPE analysis, and not at the other areas of SL5 that are unaffected in SHAPE. As seen in free SL5, the bulges serve as dynamic hinges giving flexibility to the surrounding stems. In the simulations where the cylinders started away from the RNA, they quickly localized ON those hinges, reducing flexibility of the hinge drastically (in regions W, L, N, R). From studies with three base bulges (on HIV TAR) we know that such hinges can open and from such a binding position the cylinder can reorient and insert though this can take very long on the time scales of simulations[51]; we can model this by pre-positioning the cylinder at or close to this position. Once the cylinder is in the SL5 bulge (Fig. 4A, cylinder D), the simulations show that the helical structure of the surrounding stems is disturbed, opening up the stem nucleotides to attack from 1M7, and this is confirmed experimentally in SHAPE leading to an increase in the signal in these regions (around L and M and towards W, close to the RT primer).

**Figure 4.**
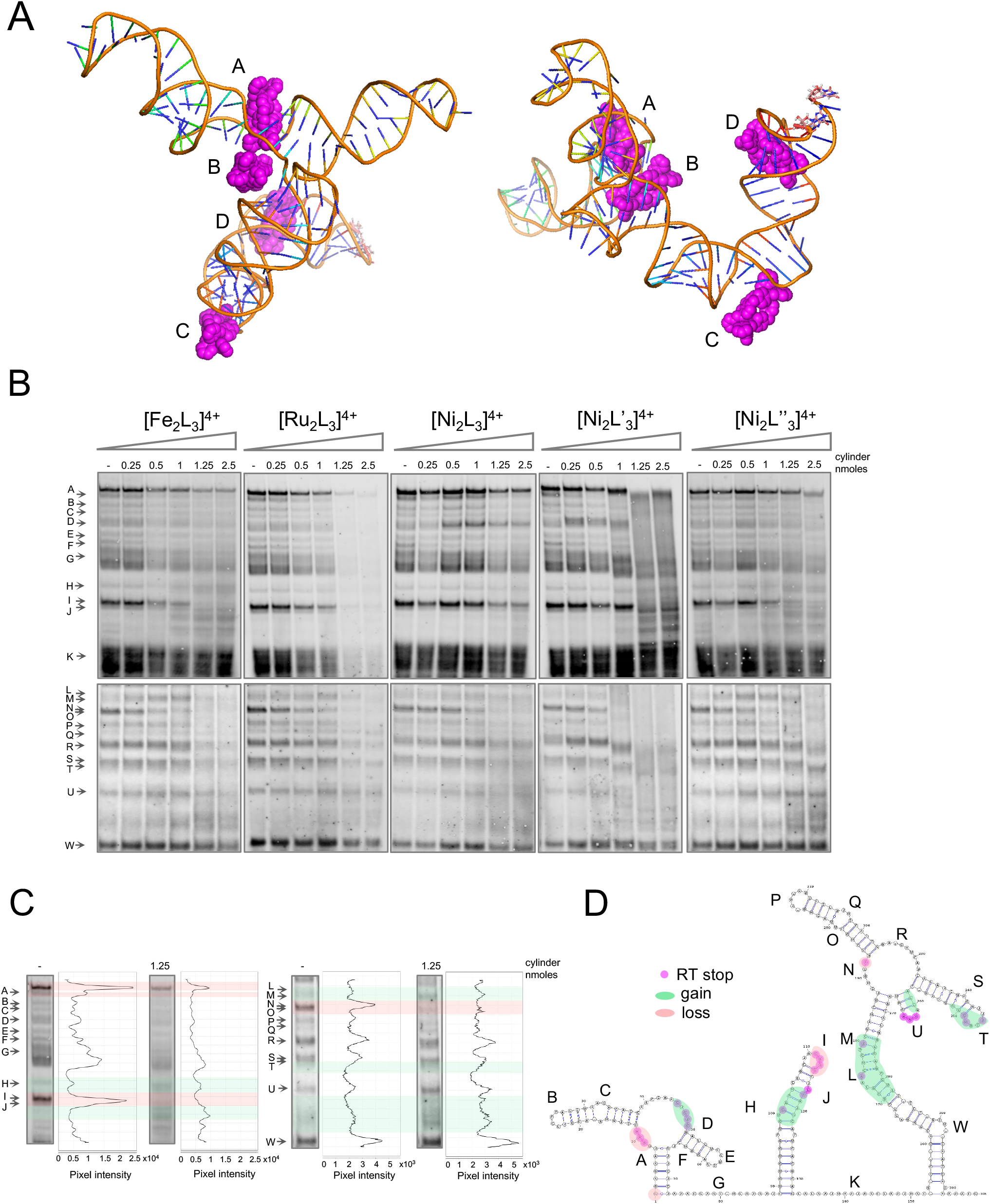
(A) View from two angles of a representative snapshot of a simulation of 4 cylinders on the SL5 RNA, revealing the same interaction points as indicated experimentally by SHAPE. Cylinder A is threaded through the central cross (4 way junction) with cylinder D threaded through the 3-base bulge at W. Cylinder B is at position N and cylinder C at position L. (B) SARS-CoV-2 5’ UTR folding in the absence (lane 1) and at increasing concentrations (lanes 2-6) of five different cylinders. Cylinders were incubated with the viral 5’ UTR (0.05nmoles) followed by SHAPE (acylation, reverse transcription and electrophoresis). (C) Band intensity of lanes 1 (without cylinder) and 5 (with) of the [Fe_2_L_3_]^4+^ gel. (D) SARS-CoV-2 5’ UTR diagram showing the RNA regions where the folding was affected by the presence of cylinder, as indicated by SHAPE.

In addition to the bulge as a site of binding, in the simulations the cylinder can also insert into the cavity at the central cross (4 way junction) (Fig. 4A, cylinder A), protecting A193. This cavity is larger than the 3-base bulge and thus although the binding site may not offer as good a structural fit, it will be kinetically quite accessible. The binding also to this site was confirmed experimentally by the disappearance of this SHAPE signal (A193, RNA position N) at increased concentration of cylinder. At the loading of cylinder used in the simulation, interaction with the stems containing regions U and T was not observed. The SHAPE results suggest that these regions are also affected as the loading increases.

In SL3 there are no large bulges similar to that found in SL5, however mismatched pairs create a distortion on the helical structure that can lead to exposure of nucleotides to 1M7. Specifically, molecular dynamics simulations (Fig. 5) on the free RNA (no cylinder) revealed short lived pairings of different types from G96:C126 to A102:U120. Furthermore, higher up the stem U104:A118 to G106:G115 is also a region of multiple cross strand pairings. Equally important for understanding the SHAPE results is the transient stacking between this stem’s nucleotides revealed in the 3D model.

**Figure 5.**
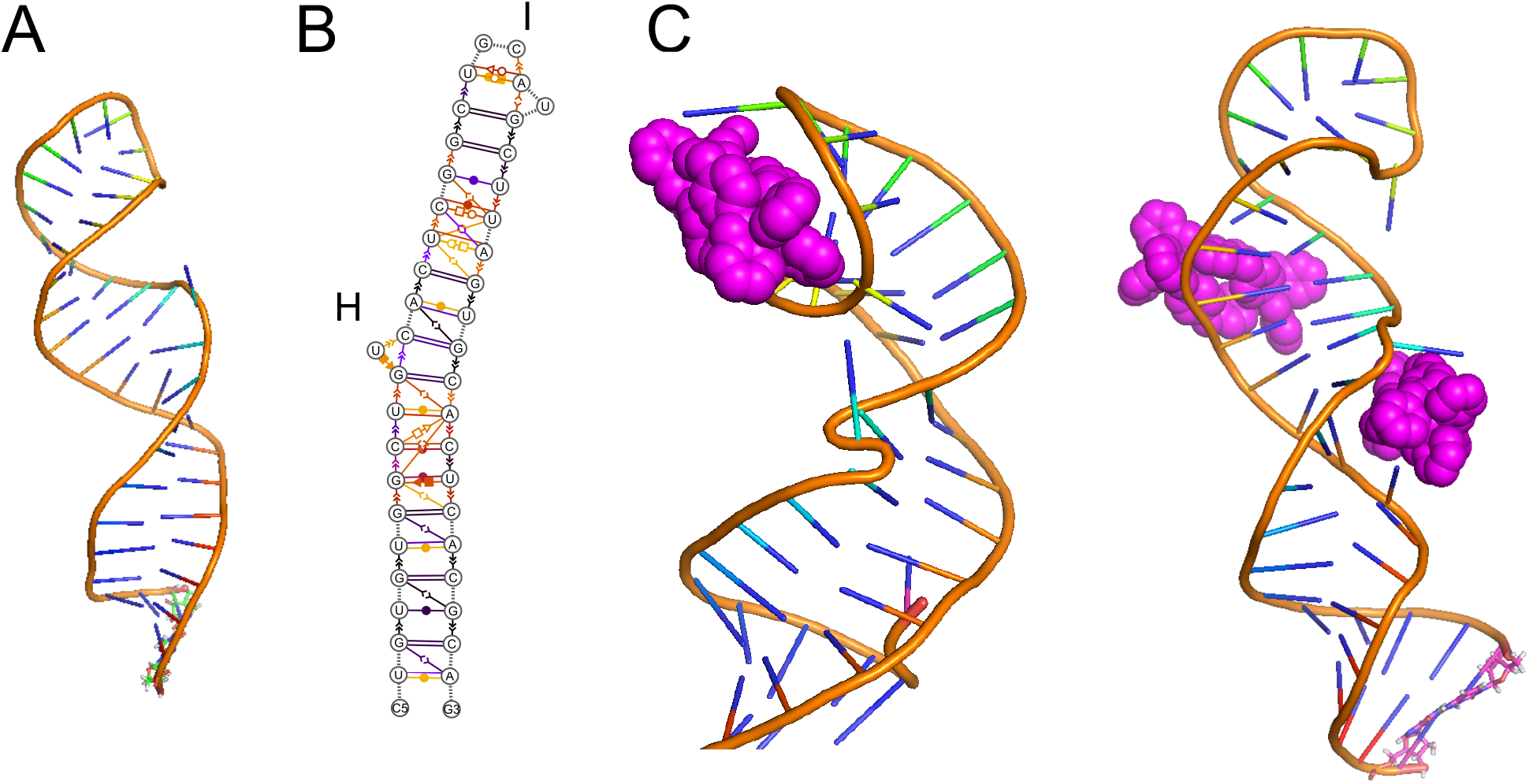
(A) Snapshot of the dynamic three-dimensional structure of the SL3 RNA from MD simulations together with a Leontis Westhoff diagram (B) highlighting the dynamic base-pairing within the structure. (C) View of representative snapshots of simulations of cylinders on the SL3 RNA, showing binding at the stem loop and on the stem as also revealed by the SHAPE analysis.

In the presence of cylinders, we observed that the cylinder is attached to the stem loop (Fig. 5C) in a stable manner, decreasing the flexibility of those residues and thus protecting the loop nucleotides from acylation, where we saw a reduced signal in SHAPE (Fig. 4B region I). Cylinders can also bind lower on the stem (region H/J) and this leads to an enhancement of acylation as seen on the stem of SL5.

Alongside the SHAPE experiments with the [M_2_L_3_]^4+^iron(II) cylinder (M=Fe), we also compared the analogous nickel(II) and ruthenium(II) cylinders (M=Ni, Ru; Fig. 4B). Changing the metal does not affect the overall cylinder structure or charge, and analogous patterns/effects are seen in the SHAPE mapping confirming that they bind the RNA at the same locations and it is the cylinder shape/charge that is responsible for the binding not the choice of metal. High cylinder excess (two last conditions, 1.25 and 2.5 nmoles corresponding to 25 and 50 cylinders per UTR) in most cases severely affected RNA structures and so SHAPE bands become less well defined indicating more random RT stops. In PCR experiments the [Ru_2_L_3_]^4+^ cylinder is stable to the heat cycles and can inhibit polymerase amplification[52]; the reverse transcription efficiency seems similarly affected at the highest concentrations of this cylinder. Some small gel shifts are also observed at high cylinder loading, possibly suggesting some cylinder-binding to the DNA transcript.

We also tested the effect of two substituted cylinders based on ligands L’ and L”, to confirm the key binding area of the cylinder design (Fig. 4B). These cylinders bear additional aryl rings at their ends while the central regions of the cylinder (which insert into the junctions/bulges) are unchanged. Both show similar patterns in the SHAPE analysis to the cylinders of ligand L, but while [Ni_2_L”_3_]^4+^ had very a similar impact on folding, the isoquinoline cylinder [Ni_2_L’_3_]^4+^ caused some changes in the SHAPE pattern even at the lowest cylinder concentrations. The results suggest that it may be possible to modify the cylinder structure to modulate the affinity for the binding sites.

Having established that the cylinder can bind and modify the structure and reactivity of the SARS-CoV-2 5’ UTR in vitro, we explored their potential to inhibit viral replication in cellulo. Simian Vero cells were infected with SARS-CoV-2 virus England 2 (Wuhan strain; identical 5’ UTR to reference sequence) in the presence or absence of the Ru and Ni cylinders, [M_2_L_3_]^4+^ (M=Ru, Ni), and the frequency of cells expressing the viral encoded Spike glycoprotein quantified (Fig. 6). Both cylinders reduced spike-expressing cells in a dose responsive manner, with the ruthenium cylinder being more effective and reducing the frequency of infected cells to <5% at the highest doses tested (75 µM). MTT cell metabolic activity/viability assays confirmed that the cylinder is not cytotoxic in the timeframe of these experiments (See Supplementary Information).

**Figure 6.**
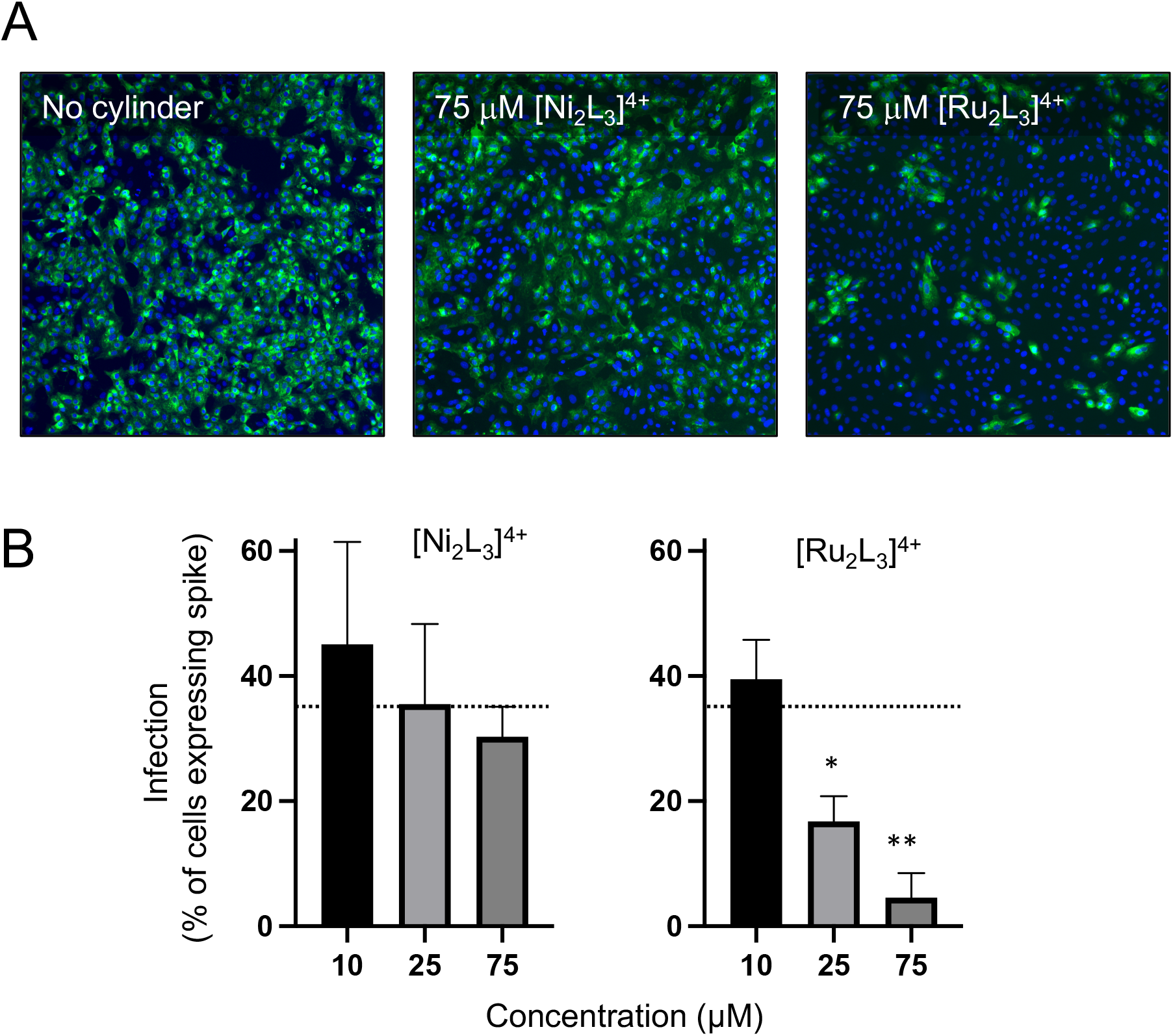
Effects of the [M_2_L_3_]^4+^ (M=Ru, Ni) cylinders on SARS-CoV-2 infection of Vero cells. Cells were infected with SARS-CoV-2 (MOI=0.04) in the presence or absence of cylinders and fixed at 48 hours post-infection and spike expression quantified by rabbit anti-spike monoclonal antibody (CR3022) and mouse anti-rabbit Alexa 555 (green). Cell nuclei were visualised with Hoechst 33342 (blue). Total cell numbers and percentage of Spike expressing cells were enumerated by high content imaging at x10 magnification using a CellInsight CX5 high content microscope (Thermo Fisher Scientific). (A) Representative images of untreated or 75 μM [Ni_2_L_3_]^4+^ or [Ru_2_L_3_]^4+^ treated cells. (B) Data represents the mean from three independent experiments and the error bars show standard deviations. Statistical analyses show Student’s t tests with Welch’s correction compared to no cylinder (dotted line), * p = 0.0168 and ** p = 0.0037.

We have shown that by combining experimental SHAPE results with Molecular Dynamics simulations we can create 3D models of the structure and dynamics of key individual stems that make up the 5’ UTR of SARS-CoV-2 These stems contain a number of intriguing structural motifs also found in the UTRs of other viruses, and which offer the possibility of developing new antiviral agents that act against a broad spectrum of diseases. The unique nucleic acid binding activity of the supramolecular cylinders is ideally suited to target these types of structures and we show that the cylinders can bind non-covalently to an RNA bulge in stem loop 5, as well as the central cross (4-way junction) of that loop. The ability to bind at different crucial RNA structural sites that are essential for virus replication limits the opportunity for the virus to mutate and to evade drug action. In line with their RNA binding, these nanosized supramolecular helicates inhibit infection at concentrations where they have negligible cellular toxicity.

These helicate cylinders are currently the only metallo-supramolecular architectures that have been demonstrated to thread through RNA bulge and junction structures, but there is a growing interest in metallo-supramolecular designs to bind nucleic acid structures.[53, 54] The results herein suggest that nucleic acid binding metallo-supramolecular architectures, and the cylinder designs in particular, merit further exploration as antiviral agents.

## Acknowledgements

This work was funded by the EPSRC Physical Sciences for Health Centre (LM, TC, JSC, MJH: EP/L016346/1), BBSRC MIBTP (NJC, CAJH, BB/M01116X/1) with Sygnature Drug Discovery (NJC, CASE 1940003), and an EU Marie Curie Fellowship (AG, H2020-MSCA-IF-2018-844145). PG and KW were supported by a Sir Henry Dale Fellowship from the Wellcome Trust and the Royal Society (200473/Z/16/Z) and ZS by a Medical Research Foundation intermediate career fellowship (MRF-169-0001-F-STAM-C0826). JAM is funded by a Wellcome Investigator Award (IA) 200838/Z/16/Z, UK Medical Research Council (MRC) project grant MR/R022011/1 and Chinese Academy of Medical Sciences (CAMS) Innovation Fund for Medical Science (CIFMS), China (grant number: 2018-I2M-2-002). We thank Henri Huppert for expert assistance with developing CellInsight quantification algorithms to measure infection in the microneutralisation assay and Jack Dismorr for synthetic assistance. Simulations used the Bluebear and Castles HPC facility (U. Birmingham).[55]

