## Supplementary Information for "Supramolecular cylinders target bulge structures in the 5’ UTR of the RNA genome of SARS-CoV-2 and inhibit viral replication"

### A. METHODS

#### 1. Software Used

***Secondary structure prediction software*** can be found at the following links;

RNA-fold http://rna.tbi.univie.ac.at/cgi-bin/RNAWebSuite/RNAfold.cgi
vfold http://rna.physics.missouri.edu/index.html
SPOT-RNA https://github.com/jaswindersingh2/SPOT-RNA
E2EFOLD https://github.com/ml4bio/e2efold
vsfold http://www.rna.it-chiba.ac.jp/~vsfold/vsfold5/
mxfold2 https://github.com/keio-bioinformatics/mxfold2
visualisation of RNA secondary structure VARNAv3-93 http://varna.lri.fr/

***3D structure prediction***
Farfar2 was used for the 3d structure prediction, summary of all elements of the UTR can be found independently at;
farfar2 - https://github.com/DasLab/FARFAR2-SARS-CoV-2
https://rosie.rosettacommons.org/farfar2
https://www.rosettacommons.org/docs/latest/FARFAR2

Comparison with other software creating 3D structures can be found here;
Vfold3D - http://rna.physics.missouri.edu/vfold3D/index.html
ifoldrna https://dokhlab.med.psu.edu/ifoldrna/
rnafold_3d http://rna.tbi.univie.ac.at/

***Sequence alignment***
MAFFT version 7 https://mafft.cbrc.jp/alignment/software/source.html

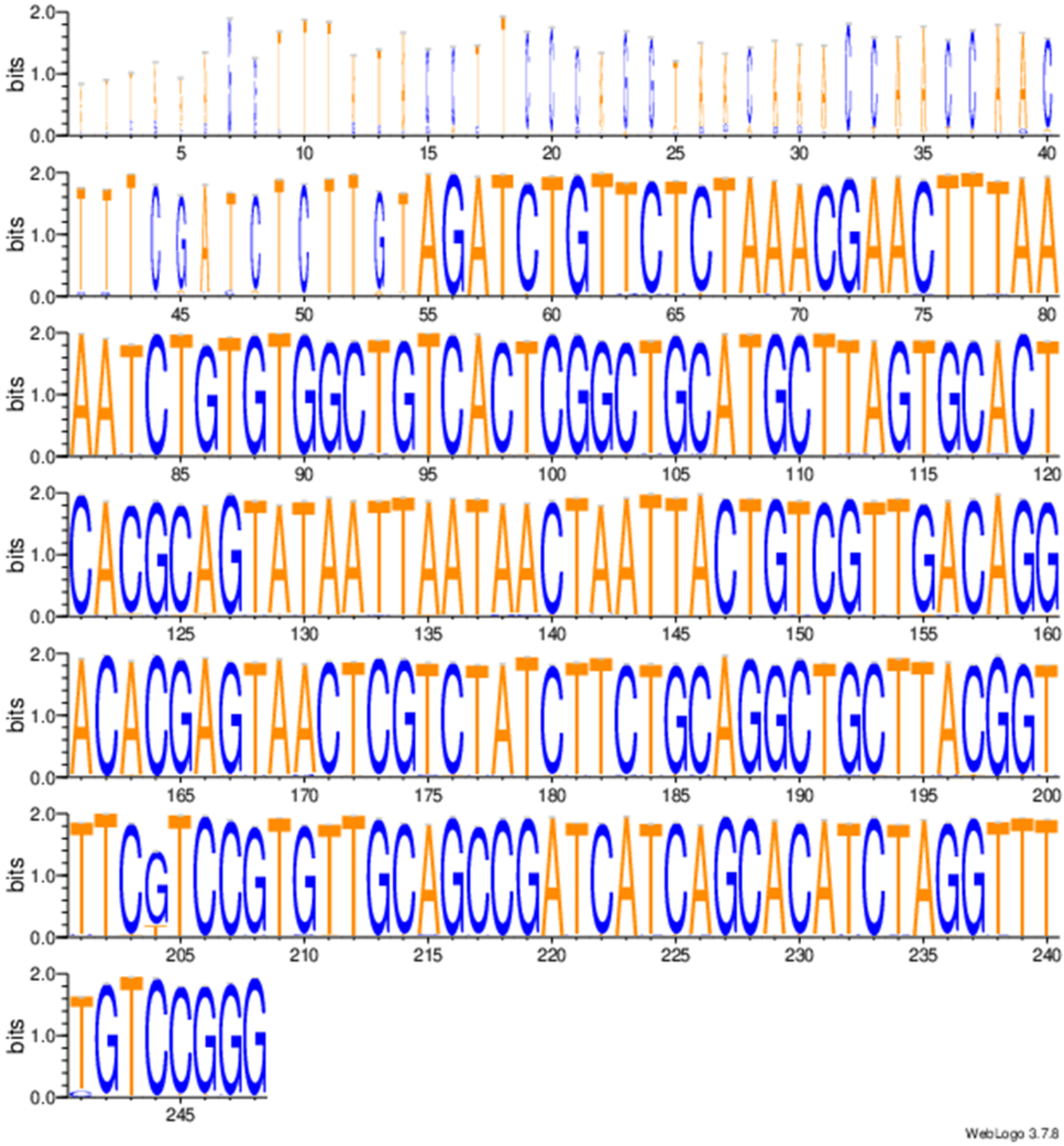

**Figure S1** *Weblogo graphical representation of sequence alignment for complete sequences from GISAID[1] referenced to NCBI Reference Sequence: NC_045512.2 on alignment with MAFFT[2]*
Raw data and analysis of secondary strucures of potential variants included in zenodo folder doi; 10.5281/zenodo.4628044

.

### 2. SHAPE RNA Analysis

#### a. SARS-CoV-2 template construction

The SARS-CoV-2 5' UTR sequence was ordered as gBlock (IDT). The UTR sequence was preceded by the T7 RNA Polymerase promoter. gBlock was flanked with XbaI and KpnI restriction sites to facilitate cloning. The template was amplified by nested PCR using primers FWD and REV. The PCR product was digested with XbaI/KpnI, gel purified and used for in vitro RNA synthesis.

full g-block
ACGCGATCGTATCTAGATAATACGACTCACTATAGGGAGAATTAAAGGTTTATACCTTCCCAGGT
AACAAACCAACCAACTTTCGATCTCTTGTAGATCTGTTCTCTAAACGAACTTTAAAATCTGTGTG
GCTGTCACTCGGCTGCATGCTTAGTGCACTCACGCAGTATAATTAATAACTAATTACTGTCGTTG
ACAGGACACGAGTAACTCGTCTATCTTCTGCAGGCTGCTTACGGTTTCGTCCGTGTTGCAGCC
GATCATCAGCACATCTAGGTTTCGTCCGGGTGTGACCGAAAGGTAAGATGGAGAGCCTTGTCC
CTGGTTTCAACGAGAAAACACACGTCCAACTCAGTTTGCCTGGTACCTGCACATAGG

5'utr 5-->3 (after XbaI/KpnI)
ATTAAAGGTTTATACCTTCCCAGGTAACAAACCAACCAACTTTCGATCTCTTGTAGATCTGTTCT
CTAAACGAACTTTAAAATCTGTGTGGCTGTCACTCGGCTGCATGCTTAGTGCACTCACGCAGTA
TAATTAATAACTAATTACTGTCGTTGACAGGACACGAGTAACTCGTCTATCTTCTGCAGGCTGCT
TACGGTTTCGTCCGTGTTGCAGCCGATCATCAGCACATCTAGGTTTCGTCCGGGTGTGACCGA
AAGGTAAGATGGAGAGCCTTGTCCCTGGTTTCAACGAGAAAACACACGTCCAACTCAGTTTGC
CT

Fwd primer: ATCGTATCTAGATAATACGACTCACTATAGG
Rev primer GTGCAGGTACCAGGCAAAC
RT1 primer CGAGTTACTCGTGTCCTGTC
RT2 primer GCAAACTGAGTTGGACGTG

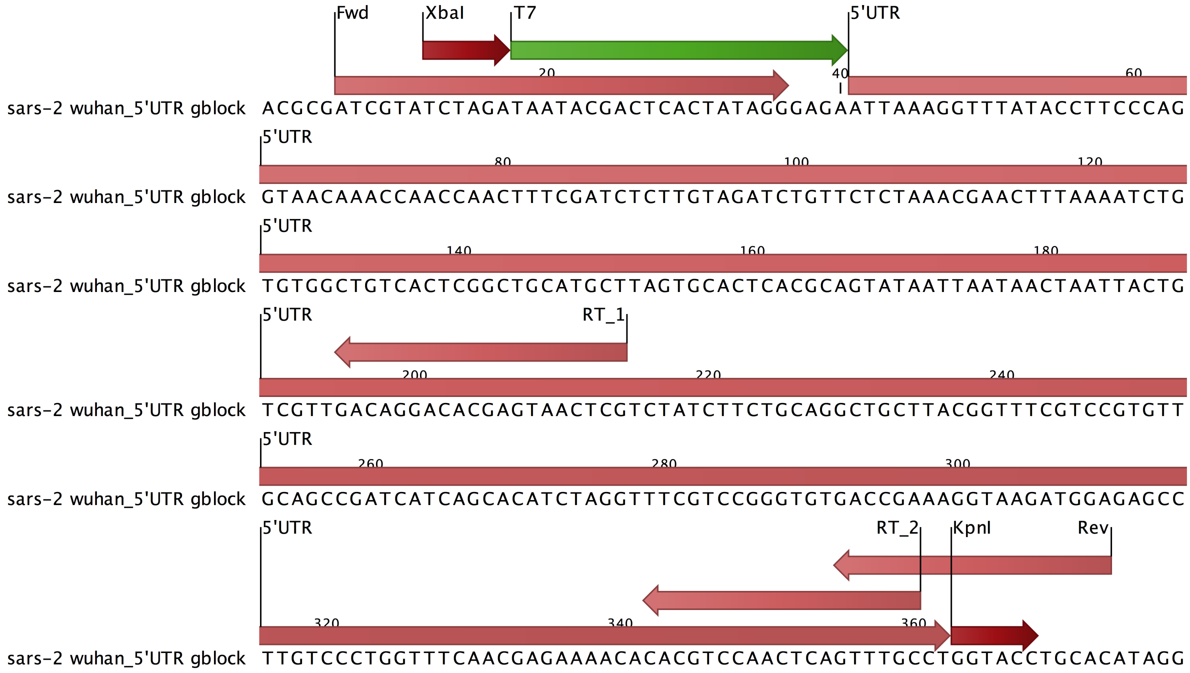


**Figure S2:** *SARS-CoV-2 template construction*

#### b. In vitro RNA synthesis

T7 MEGAscript (Ambion) was used. 200vng of DNA template was mixed with NTPs and T7 RNA polymerase in final volume 30vμl volume and incubated 3 h at 37˚C. The template was removed by TurboDNase (10U at 37˚C for 10 minutes) and precipitated with EtOH/ammonium acetate to remove unincorporated nucleotides. Precipitated RNA was suspended in 30 μl RNase-free water. RNA quality was checked on agarose denaturing gel. RNA concentration was measured on nanodrop and adjusted to the final concentration of 2.5 μg/μl.

**c. RNA folding**[1]

For each condition, 50pmoles (5μg) in 25μl of RNAse-free water was mixed with 25μl of 2x KAC folding buffer (final concentration 20mM Hepes pH 7.5 200mM KOAc pH 8 2mM $MgCl_2$) in a low-bind 1.5 ml tube, incubated at 75˚C for 5 min, slowly cooled down to 40˚C. Next, the sample was briefly centrifuged, placed on ice and immediately used for downstream analysis.

**d. Cylinder treatment**

All cylinders were prepared as 1mM solutions in water, and further diluted to 100μM working stocks. 50μl of folded RNA (50 pmoles) were mixed with the corresponding amount of 100μM cylinder solution and adjusted with water to a total volume of 75μl. E.g. for RNA/cylinder 1:5 ratio (5 cylinder equivalents per RNA stand), 2.5 μl (250 pmoles) of 100 μM cylinder and 22.5 μl water were added while for RNA/cylinder 1:50 (50 cylinder equivalents) ratio, 25ul (2.5nmoles) cylinder solution was added. Next 20 μl of 2x KAC folding buffer was added. The samples were moved from ice to 37C for 15 min and used directly for labeling.

**e. Labeling of flexible RNA nucleotides with 1M7**[1]

1M7 (1-Methyl-7-nitroisatoic anhydride) was prepared as 120nM stock in DMSO. 5μl of 1M7 was added (6nM final concentration) to 95 μl RNA/cylinder solution and incubated for 30min at 37˚C. Next RNA was precipitated with 400 μl EtOH, 5 μl 3 M NaOAc and 1 μl glycogen. Recovered RNA was then resuspended in 50 μl of nuclease-free water.

**f. Primer extension with IRD700-labelled primer**[2]

5 μl of labeled RNA (~500ng) was mixed with 1 μl of 10 μM 5'-IRD700 primer, 1 μl SS buffer and 5μl water. The sample was incubated for 5 min at 75˚C, slowly cooled down to 60˚C and placed on ice for 5 min. Next, 4 μl of reverse transcriptase buffer, 1 μl of 100mM DTT, 1 μl of 10mM dNTPs, 0.5 μl of RiboLock, 0.75 μl of water and 0.75 μl of SuperScript III (20 μl total volume) was added on ice. Next reverse transcription was performed at 55˚C for 45 min in a thermocycler. The reaction was stopped by adding 20 μl of 7M urea and directly used for electrophoresis.

**g. Electrophoresis in acrylamide gel 6%**

10 μl of RT/urea mixture was loaded on 20 cm long 6% acrylamide (19:1), 7M urea, 1x TBE gel and run at 13W for 2 hr. The gel was transferred onto whatmann paper and fluorescent RT products were visualised using the LI-COR Odyssey platform.

**h. Sanger sequencing**[2]

Sanger sequencing was performed on non-labeled RNA. For each reaction, 500 ng was mixed with 1 μl of 10 μM 5'-IRD700 primer, 1 μl SS buffer and 1 μl water. The sample was incubated for 5 min at 75˚C, slowly cooled down to 60˚C and placed on ice for 5 min. The mixture was added to tubes containing 4 μl of either G, A, T or C ddNTP (final concentration 2 mM) following adding of 4 μl of reverse transcriptase buffer, 1 μl of 100 mM DTT, 1 μl of 10 mM dNTPs, 0.5 μl of RiboLock, 0.75 μl of water and 0.75 μl of SuperScript III (20 μl total volume). Next reverse transcription was performed at 55˚C for 45 min in a thermocycler. The reaction was stopped by adding 20 μl of 7 M urea and directly used for electrophoresis. 10 μl of RT/urea sequencing reaction was loaded next to RT reaction on labeled RNA on 60 cm long sequencing 6% acrylamide (19:1), 7 M urea, 1x TBE gel and run at 40W for 3 hr. The gel was transferred onto whatmann paper and fluorescent RT products were visualised using the LI-COR Odyssey platform.

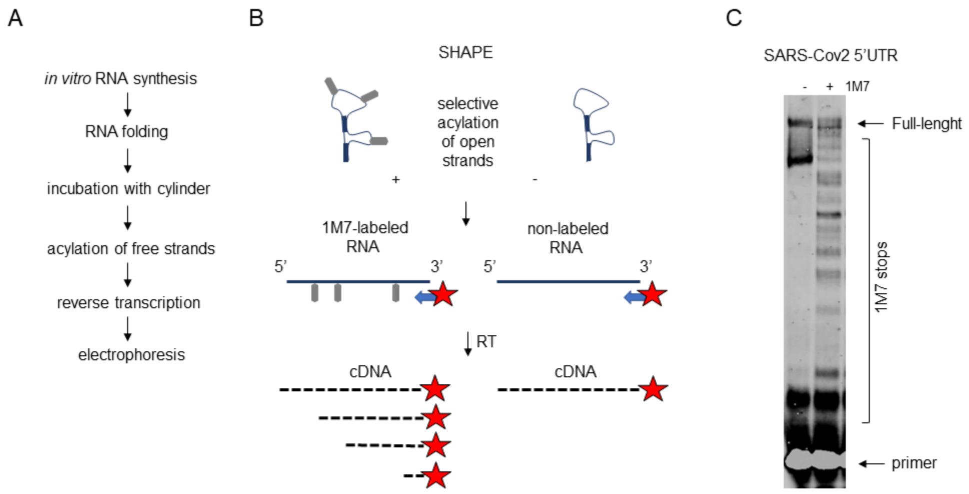


**Figure S3**. (A) An overview of experimental design. (B) SHAPE principles. (C) A comparison of RT reaction (RT2) using non-modified (-) and acetylated (+) templates.

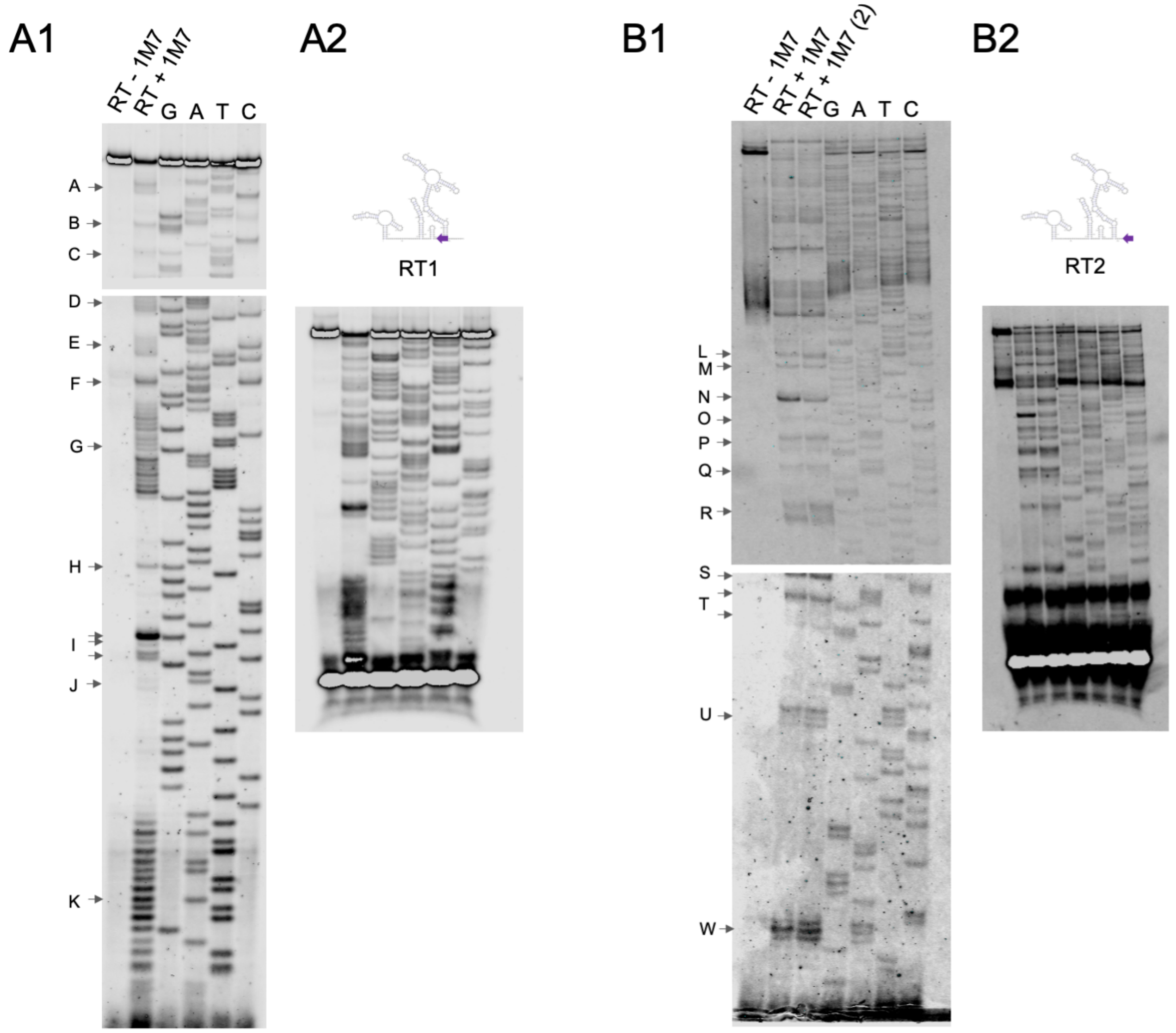


**Figure S4a**. Sanger sequencing of RT products. (A1) sequencing from RT1. Sequencing reactions were loaded along RT performed on a non-acylated and acylated template. The gel was cut to fit into the scanner. (A2) the same sequencing reaction run on a shorter gel. (B) as for (A) but sequencing performed with RT2. Moreover, an additional RT reaction was run (RT + 1M7 (2)) where the template was treated with the Fe cylinder.

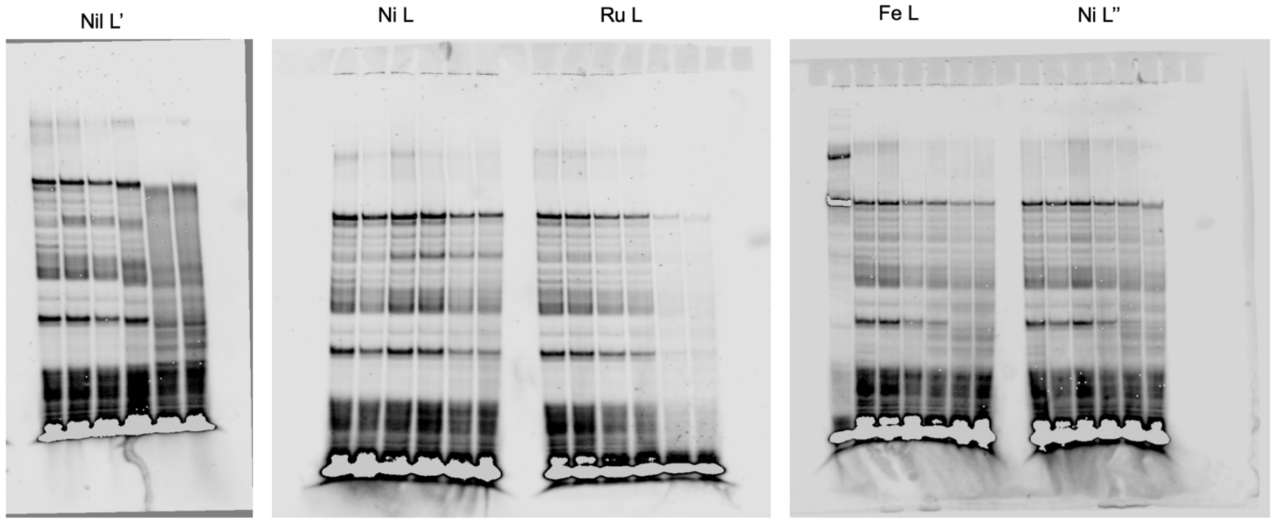


**Figure S4b.** Raw pictures of RT1

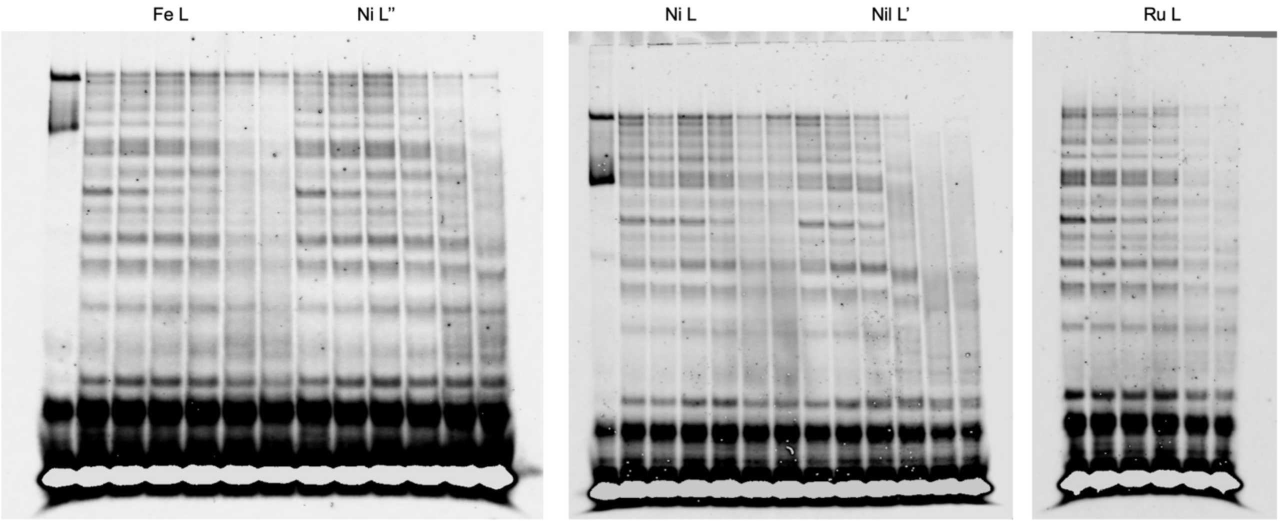


**Figure S4c.** Raw pictures of RT2

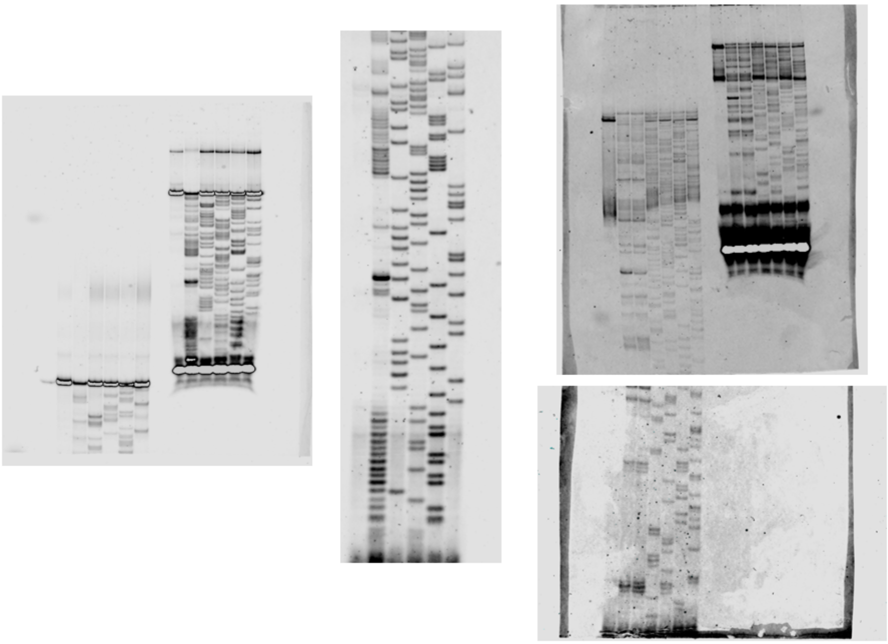

**Figure S5.** Raw pictures of sequencing

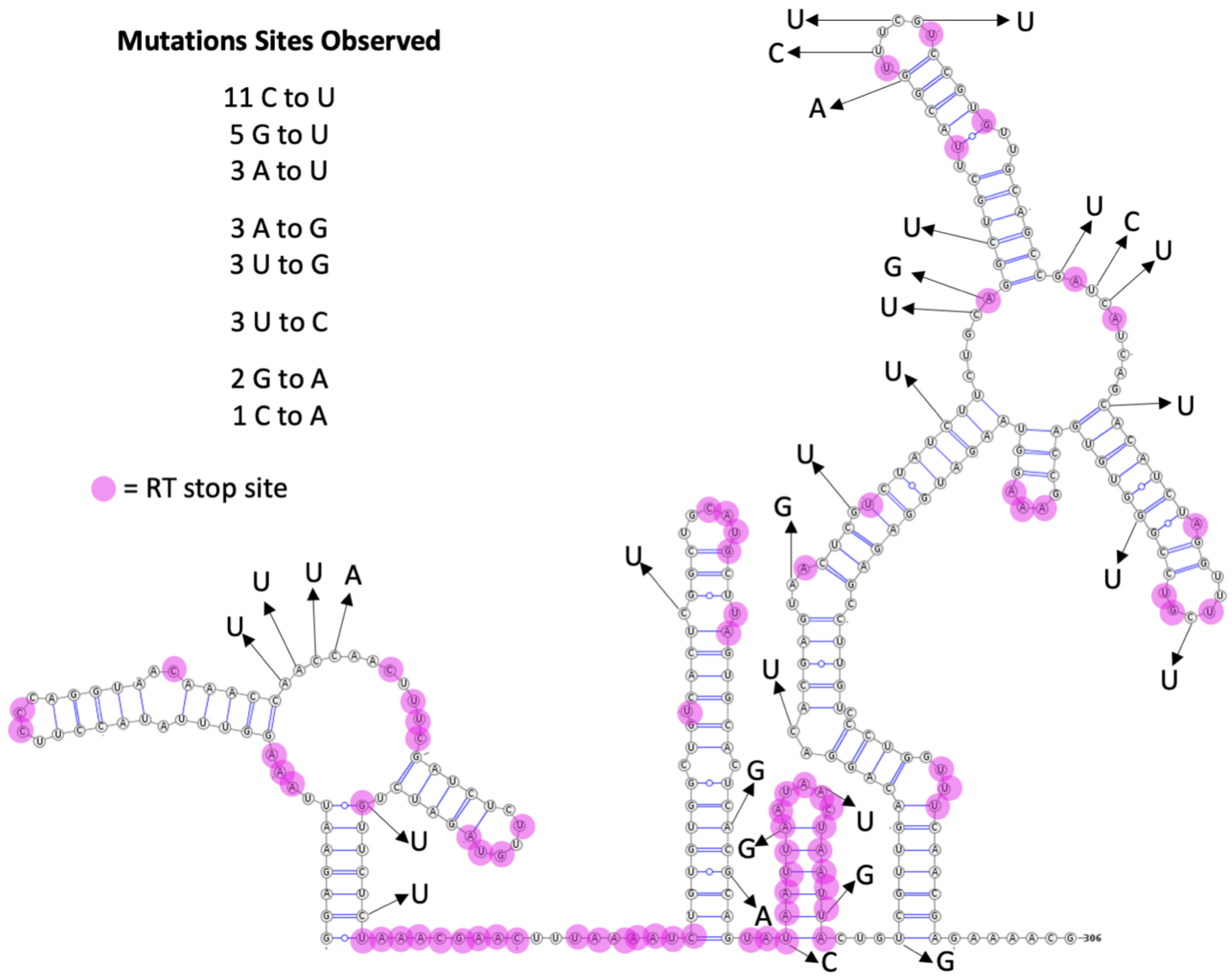


**Figure S6.** Overlay of the RT stops (main text Fig 3B) and the mutation sites (main text Fig 3B) observed in the 5' UTR of the SARS-Cov-2 viral genome, showing their proximities.

#### 3. MTT assay to assess cytotoxicity of cylinders in Vero cells

Vero cells (ATCC® CCL-81) were cultured in Dulbecco's Modified Eagle Medium (DMEM) supplemented with 10 % FBS, 1 % penicillin-streptomycin, 1 % *L*-glutamine and 1 % non-essential amino acids (Gibco). Cells were seeded into 96-well tissue culture plates (Greiner Bio-One) at 5 x $10^3$ cells/well and exposed over 48 h to either the isoquinoline or nickel/ruthenium cylinders titrated at 75 – 2 microM, vehicle control or positive cytotoxicity control. Vero cells were incubated with the tetrazolium dye 3-(4,5-dimethylthiazol-2-yl)-2,5-diphenyltetrazolium bromide (MTT) for 3 h and the absorbance of the formazan product measured at 570 nm using a multimode microplate reader (Tecan Infinite Pro 200, Männedorf, Switzerland) and cytotoxicity calculated as the number of viable cells compared to assays controls.

**A**

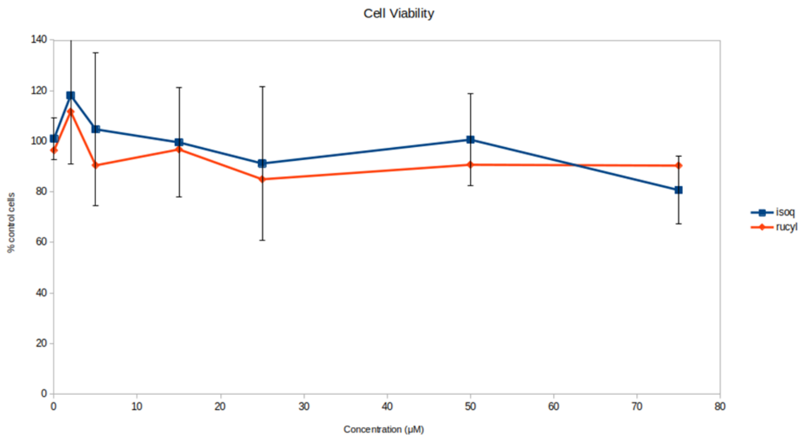


**B**

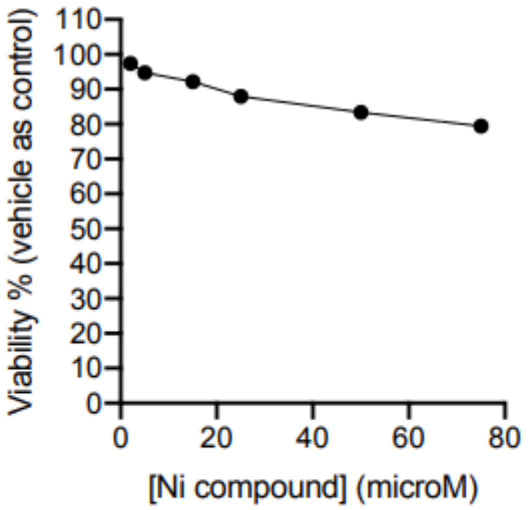


**Figure S7** A) Cell viability across concentrations of $[Ru_2L_3]Cl_4$ (red line) and $[Ni_2L'_3]Cl_4$ (blue line) cylinders derived by MTT assay B) Cell viability across concentrations of $[Ni_2L_3]Cl_4$

### 4. SARS-COV-2 viral infection assays

**Cell culture.** Vero cells were cultured in DMEM supplemented with 10% foetal bovine serum, 2mM L-glutamine, 100 U/ml penicillin, 10 µg/ml streptomycin and 1% non-essential amino acids (culture media). The cells were maintained at 37°C and 5% $CO_2$.

**Virus.** SARS-CoV-2 Strain England 2/2020, a clinical isolate sampled from a 23-year-old male during acute illness in January 2020 (GSAID Accession ID EPI_ISL_407073), was provided by Dr Christine Bruce, PHE at Porton Down. The stock was prepared by infecting 95 % confluent Vero E6 cells with virus to MOI of 0.005 and virus was harvested after 6 days. The titre was determined to be $7.0 \times 10^5$ pfu/mL by plaque assay (doi: 10.1016/j.jviromet.2021.114087).

**Infection and spike detection by immunofluorescence.** Vero cells were seeded into 96-well imaging plates (Greiner) at a density of $10^4$/well in culture media and infected after 24 hours with SARS-CoV-2 England 2 (MOI=0.04), in the presence or absence of cylinders as indicated. Infected cells were fixed in ice-cold methanol 48 hours after infection. Cells were washed in PBS and stained with rabbit anti-SARS-COV-2 spike protein, subunit 1 (CR3022, The Native Antigen Company), followed by Alexa Fluor 555-conjugated goat anti-rabbit IgG secondary antibody (Invitrogen, Thermofisher). Cell nuclei were visualised with Hoechst 33342 (Thermofisher). Cells were washed with PBS, imaged and analysed using a ThermoScientific CellInsight CX5 High-Content Screening (HCS) platform. Infected cells were scored by perinuclear fluorescence above a set threshold determined by positive (untreated) and negative (uninfected) controls.

### 5. Cylinder Synthesis

Cylinders were synthesised according to the previously published procedures[3][4][5][6][7] and isolated as their chloride salts

### 6. Molecular Dynamics Simulations

#### a. Parametrisation of supramolecular cylinder.

The crystal structure of the $[Fe_2L_3]^{4+}$ cylinder was split into 5 residues (3 ligands and 2 metal ions) that were fed to MCPB.py[8] to generated parameters for the metal centres at the wB97XD/6-31G* level of theory using Gaussian09 as well as partial charges using RESP[8]. The coordinate and parameter files were converted to GROMACS[9] using Parmed https://github.com/ParmEd/ParmEd

#### b. RNA and RNA-cylinder simulations.

Preparation of the RNA used the ROC forcefield[10] and was done using tleap[11] and the files provided. The parameters and coordinates were then converted to gromacs using Parmed.

In all systems, unless otherwise stated, the RNA was placed in a dodecahedral box with edges at least 1.2 nm from the solute, filled with TIP3P water. Initial minimisation was carried to at least 500 kJ/mol/nm or 50000 steps followed by heating and NVT equilibration for 1000ps using V-rescale modified Berendsen thermostat, coupling the cylinder with the RNA at 310K. All simulations use 2 fs time step and Parrinello-Rahman pressure coupling and PME electrostatics at 1.0nm cut-off[12].

After completion, the compressed trajectories were analysed to remove periodic boundary conditions and rotations using gromacs' trjconv program. After removing the water the trajectories were analysed with pyemma2.6[13], barnaba[14] and x3dna-dssr[15].

#### c. Pyemma analysis[13]

Different features were chosen to capture the kinetic variance that occurred during the simulations: (i) Position of centre of mass (COM) of each residue is a low dimensional and relatively efficient way to capture different states, especially including the cylinder. (ii) Taking advantage of the fact that each residue has an atom named N3, which is away from the backbone, one can easily create matrix of distances between these N3 atoms, which although high in dimensionality it captures nearly all the kinetic variance.

For each simulation, Principle Component Analysis (PCA) was carried out and the projections between the first 4 PCs plotted, followed by TiCA analysis for lag times 1 to 5000 steps. The lag time for which the fewer number of TiCA dimensions are necessary to capture 95% of the kinetic variance was chosen for further analysis. The number of clusters was chosen by examining the convergence with regards to VAMP2. Lag times for MSM model are chosen from the convergence at timescales of identified processes. Only models that use all of the states and can pass the Chapman-Kolmogorov test can continue to Perron cluster cluster analysis (PCCA) which leads to extraction of states with certain probability and structure in pdb format. Not all simulations were long enough to produce an appropriate Markov state model, and it should be noted that the Markov state models are meant to describe or sum up the particular simulations and not the whole system.

#### d. Barnaba analysis[14]

All long production molecular dynamics runs as well as states identified by PCCA were analysed using barnaba resulting in 2D Leontis/Westhof classification[16] of base interactions.

All analysis of MD runs is included in the zenodo folder: doi; 10.5281/zenodo.4628044

### B SUPPLEMENTARY RESULTS

#### 1. Secondary structure Predictions

##### *a. SPOT-RNA prediction*

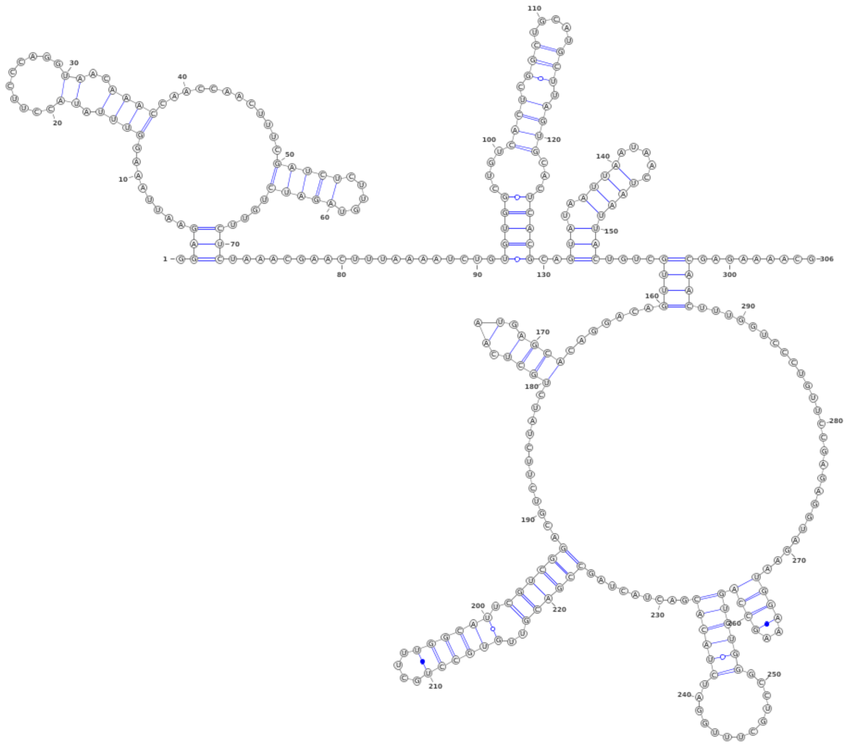

**Figure S8.** Secondary Structure prediction of the experimental RNA sequence using SPOT-RNA

##### *b. RNAFOLD prediction*

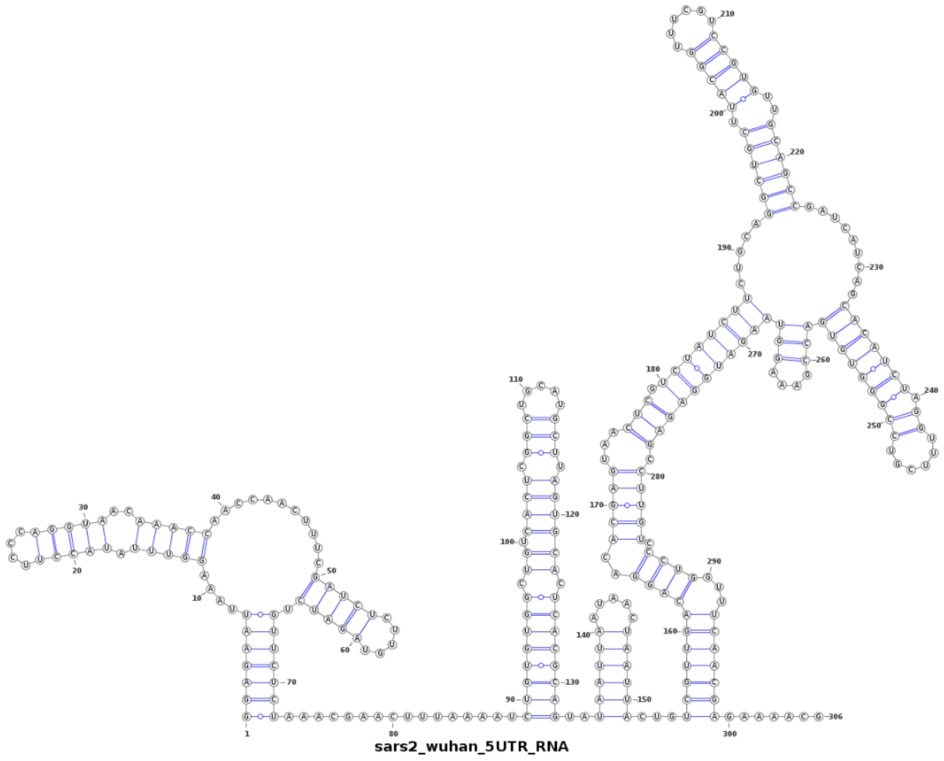


**Figure S9.** Secondary structure prediction of the experimental RNA sequence using RNAfold

*c. vfold prediction*

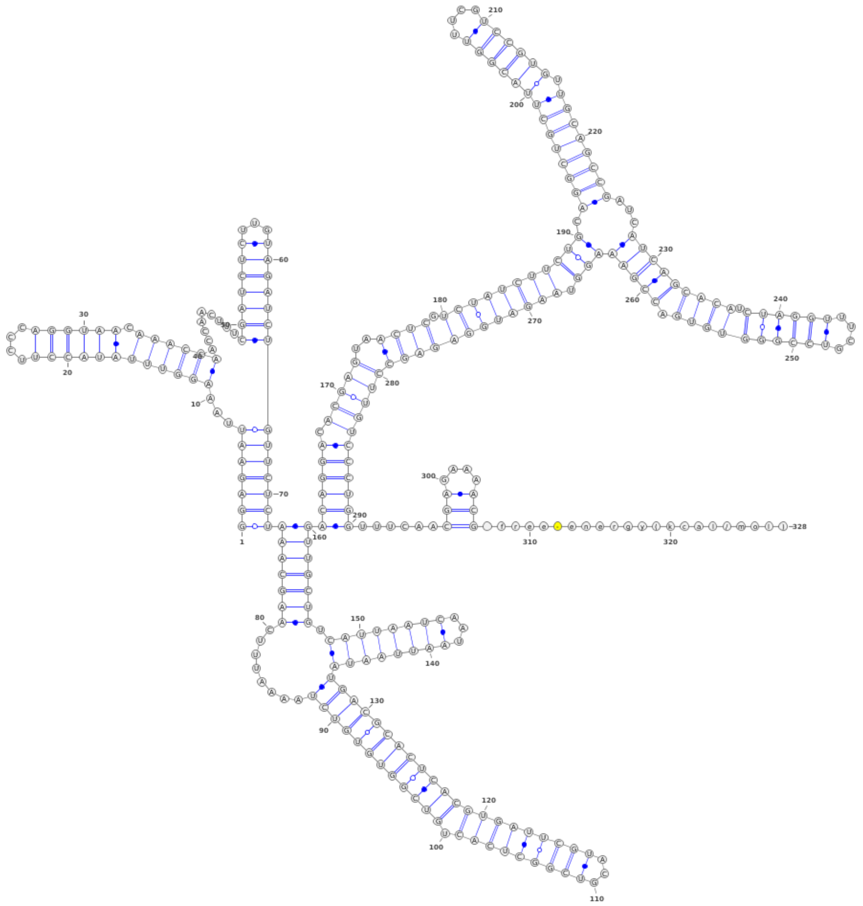

**Figure S10***: Secondary structure prediction of the experimental RNA sequence using vfold*

*d. mxfold2 prediction*

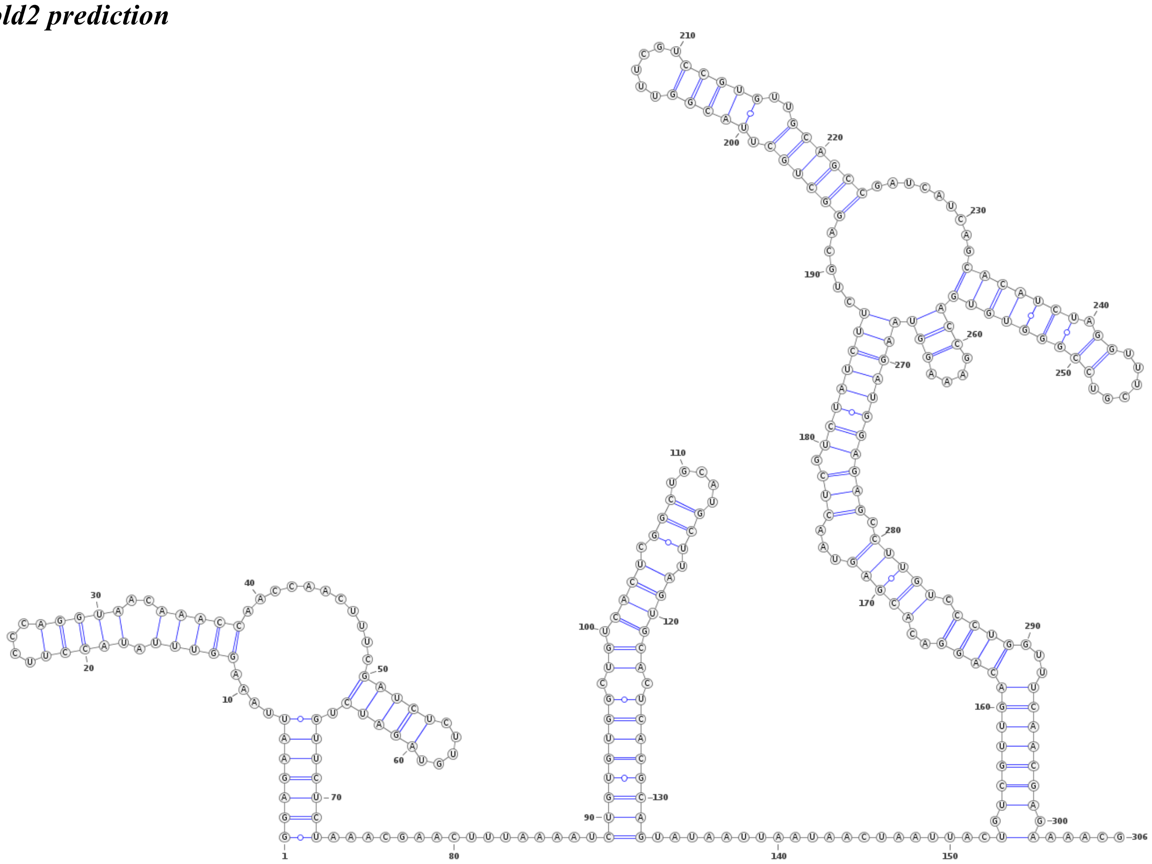

**Figure S11** *Secondary structure prediction of the experimental RNA sequence using mxfold2*

**2. 3D structure predictions compared as suitable starting points for MD simulations of SL5**

**a. RNAfold**

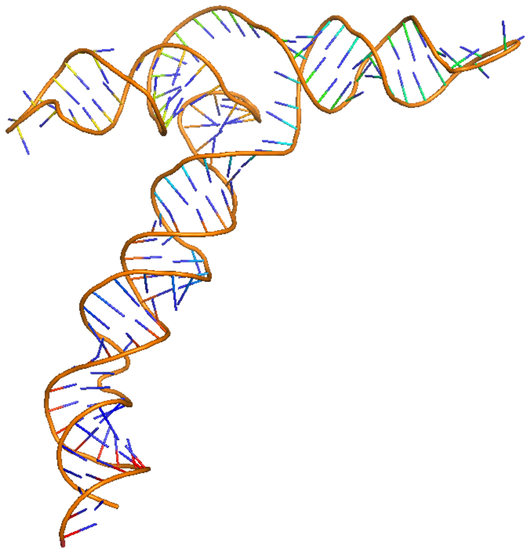

**Figure S12**. 3D structure prediction based on RNAfold secondary structure prediction of SL5

**b. FARFAR2**

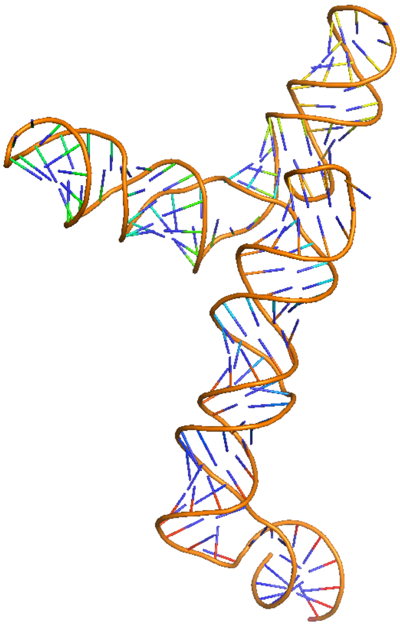

**Figure S13**. 3D structure prediction based on FARFAR2 protocol prediction of SL5

### 3. Molecular dynamics simulations of SL5

#### a. SL5 RNA (no cylinders)

combined analysis of 3 simulations of free SL5

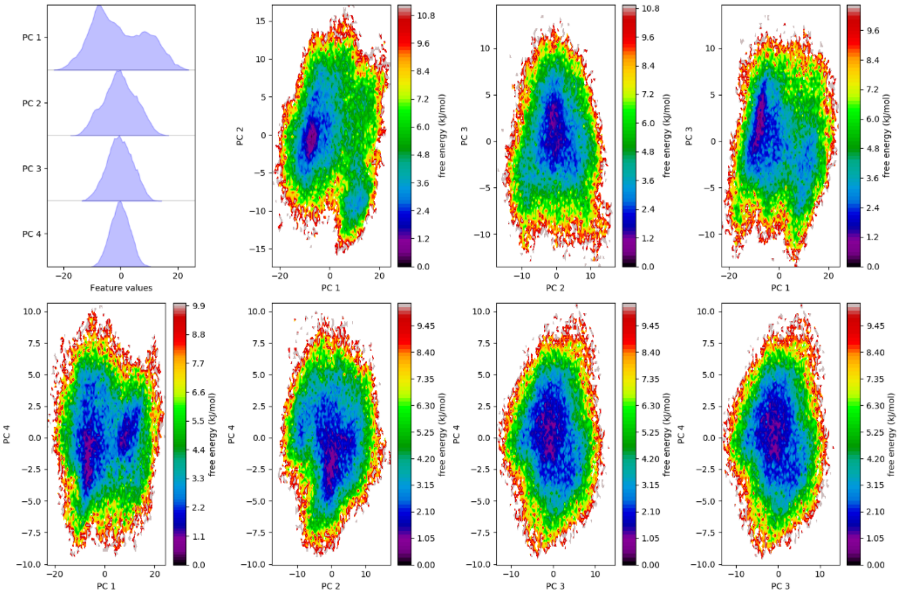


**Figure S14.** PCA of combined simulations of free sl5 simulation

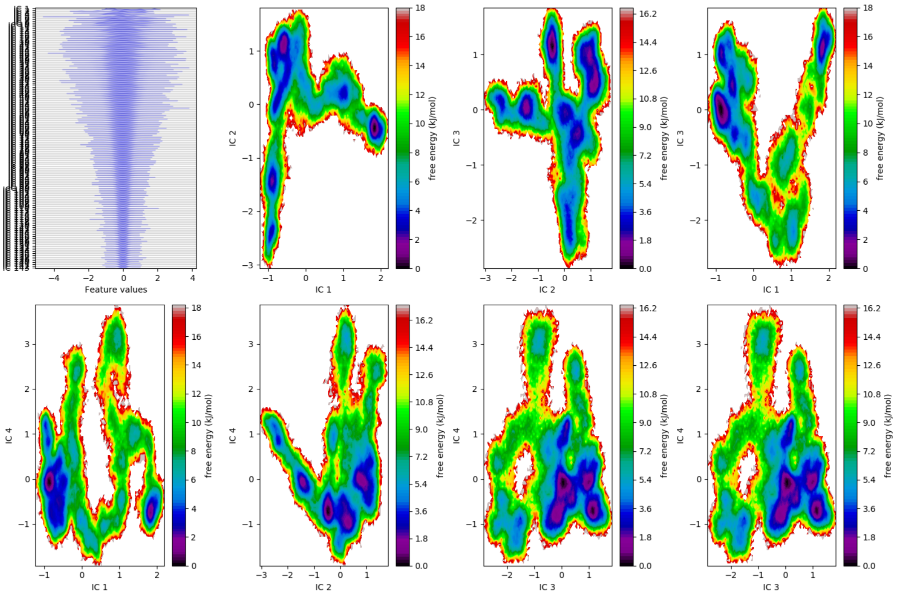


**Figure S15.** TICA of combined simulations of free sl5 simulation

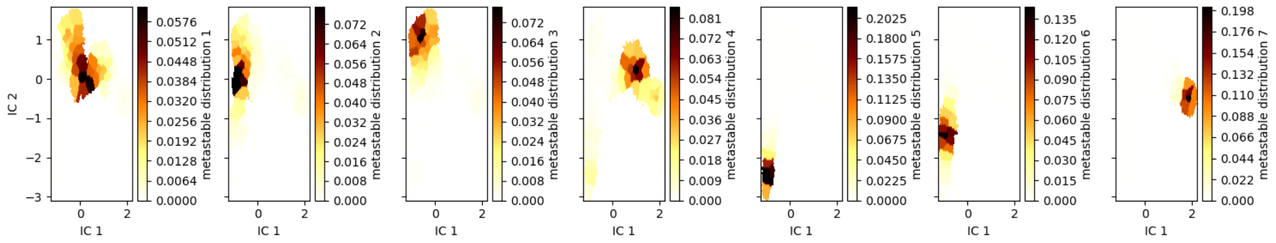


**Figure S16.** Separation of metastable states from the TICA analysis

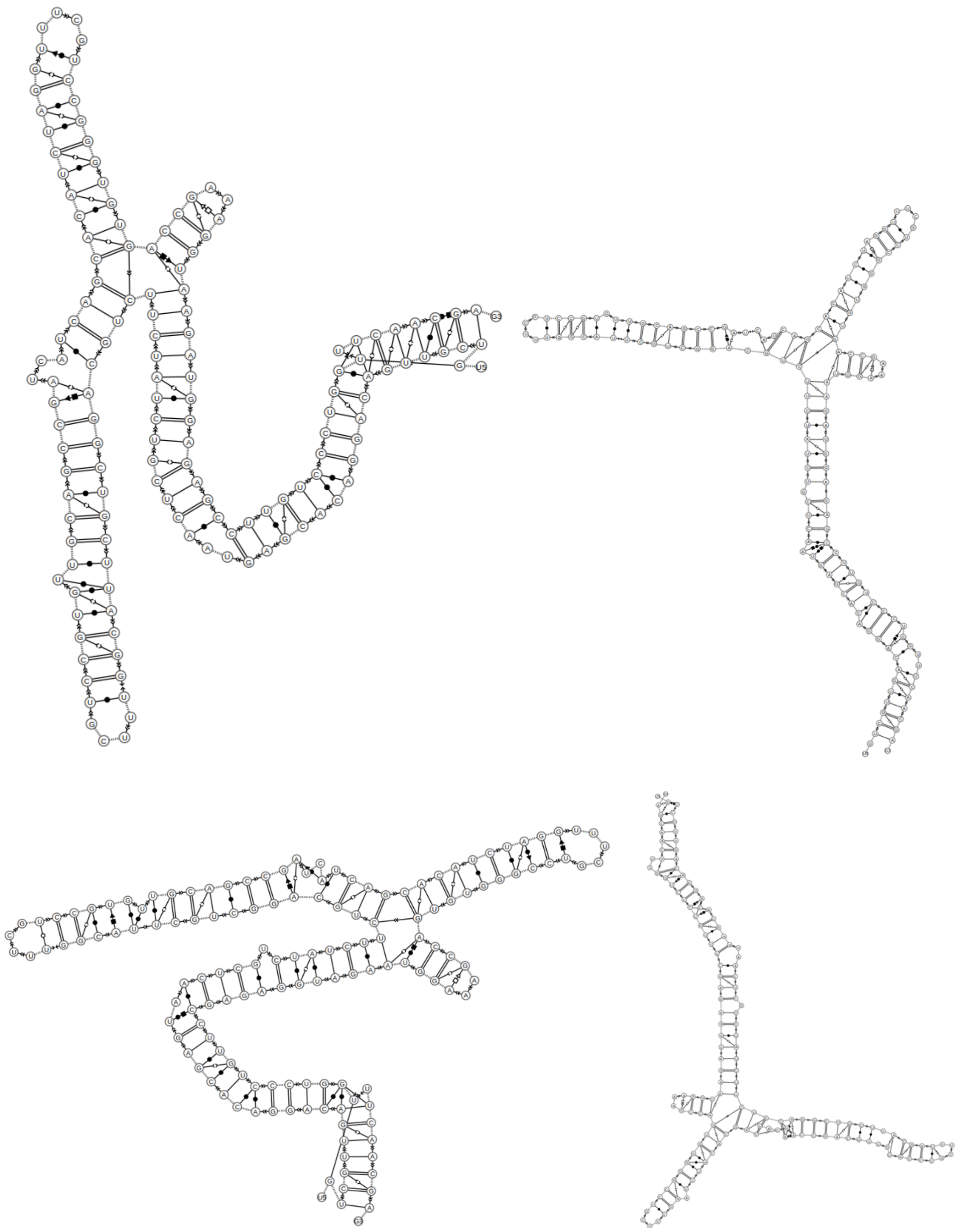

**Figure S17.** Barnaba representation of the metastable states 1, 2, 4, 6 identified in the TICA analysis of the combined simulations

**b. SL5 plus four M enantiomer cylinders introduced outside the RNA**

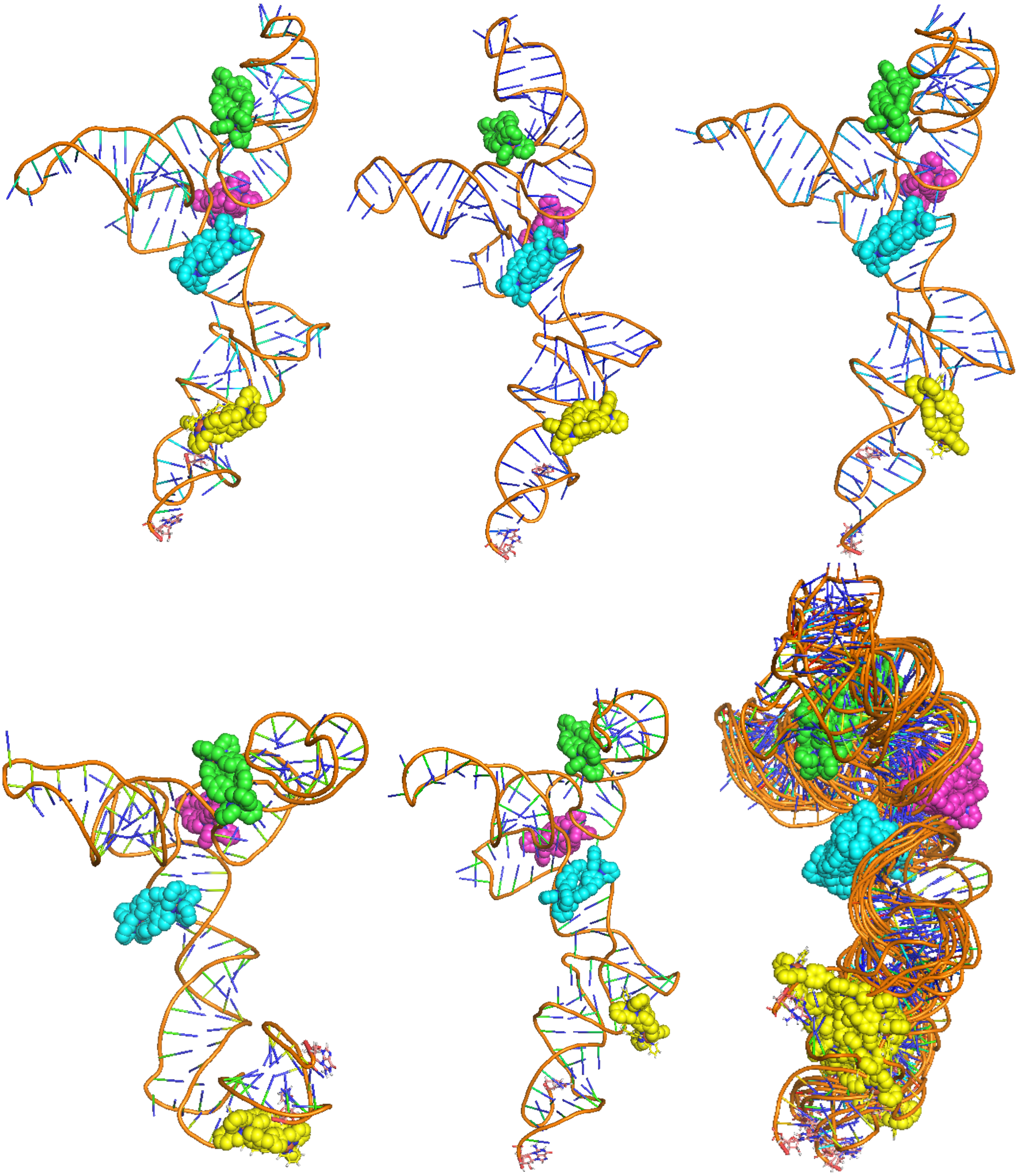

**Figure S18.** 3D structures of metastable states observed for M enantiomers on SL5, and a superposition of those metastable states; cylinders localise at both the 4WJ and the bulge site.

**c. SL5 plus M enantiomers starting with cylinders placed at the 4WJ and the bulge sites**

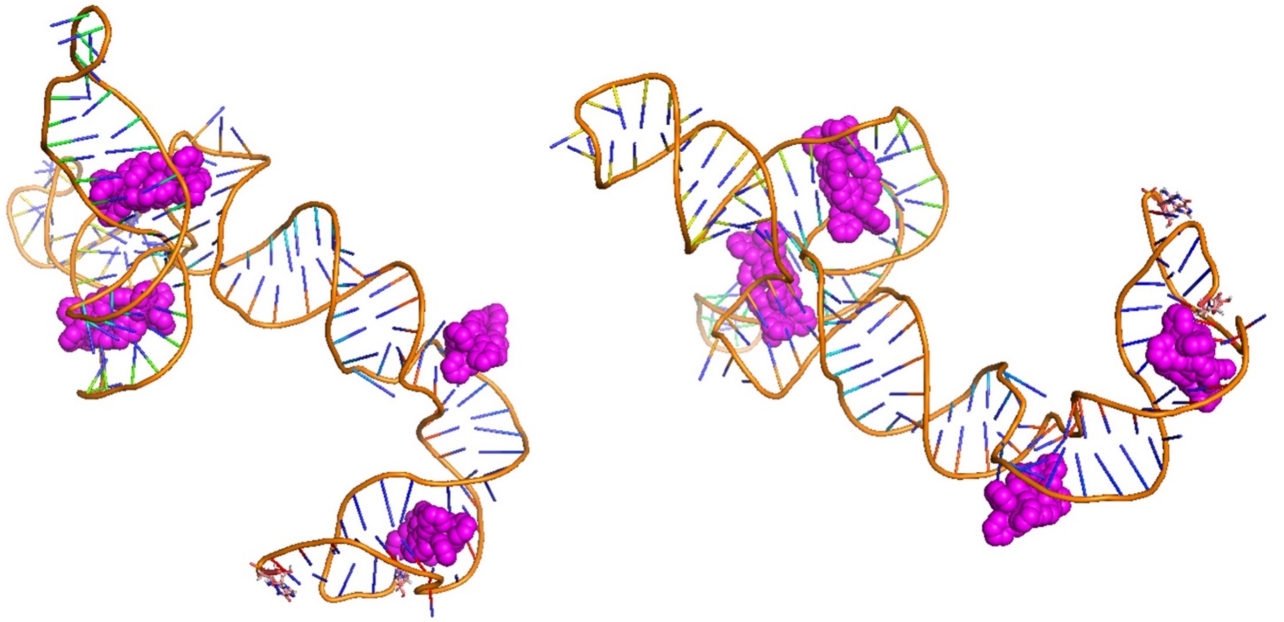

**Figure S19.** 3D structures of M enantiomer cylinders placed in bulge and junction at the outset of the simulation

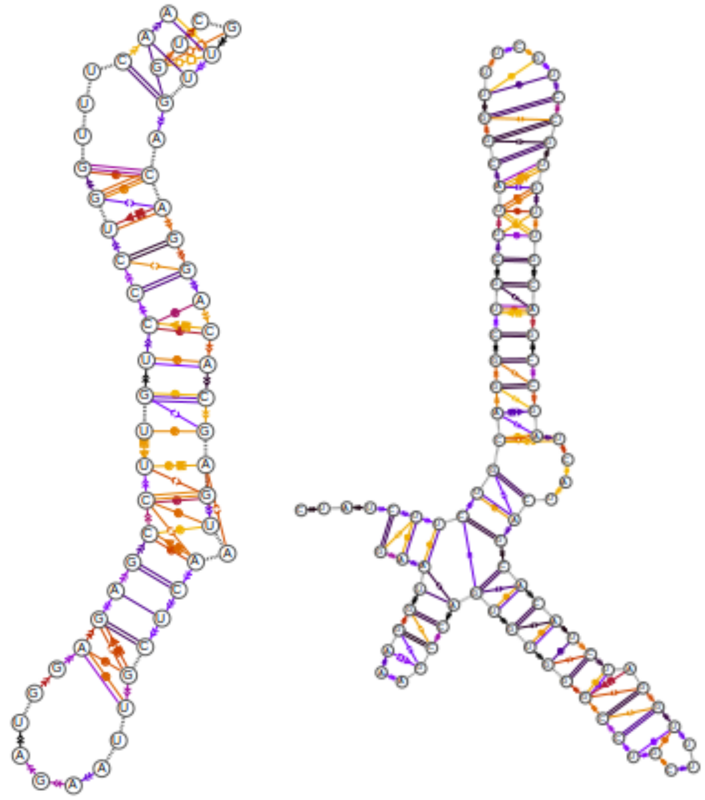

**Figure S20.** Barnaba representations of stem (A) and junction, sl5a, sl5b, sl5c (B) of M cylinder inserted simulation

**d. SL5 plus four P enantiomer cylinders introduced outside the RNA**

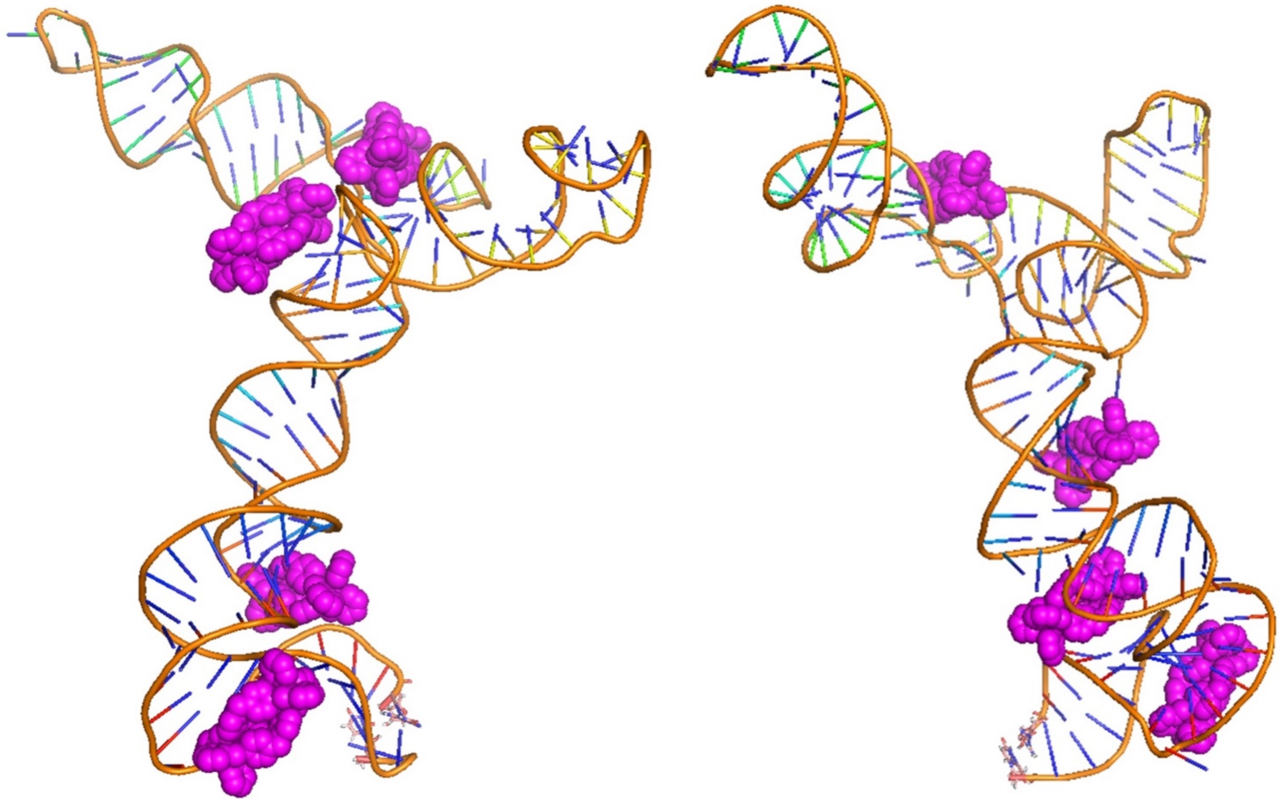

**Figure S21.** 3D structures showing P cylinders on SL5 – the cylinders are observed in very similar locations to the M enantiomers, and again localise at the 4WJ and the bulge site.

**e. SL5 plus P enantiomers starting with cylinders placed at the 4WJ and the bulge sites**

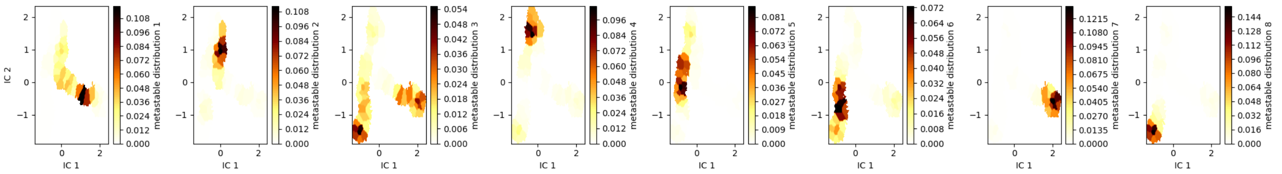


**Figure S22.** Metastable state separation of the TICA landscape

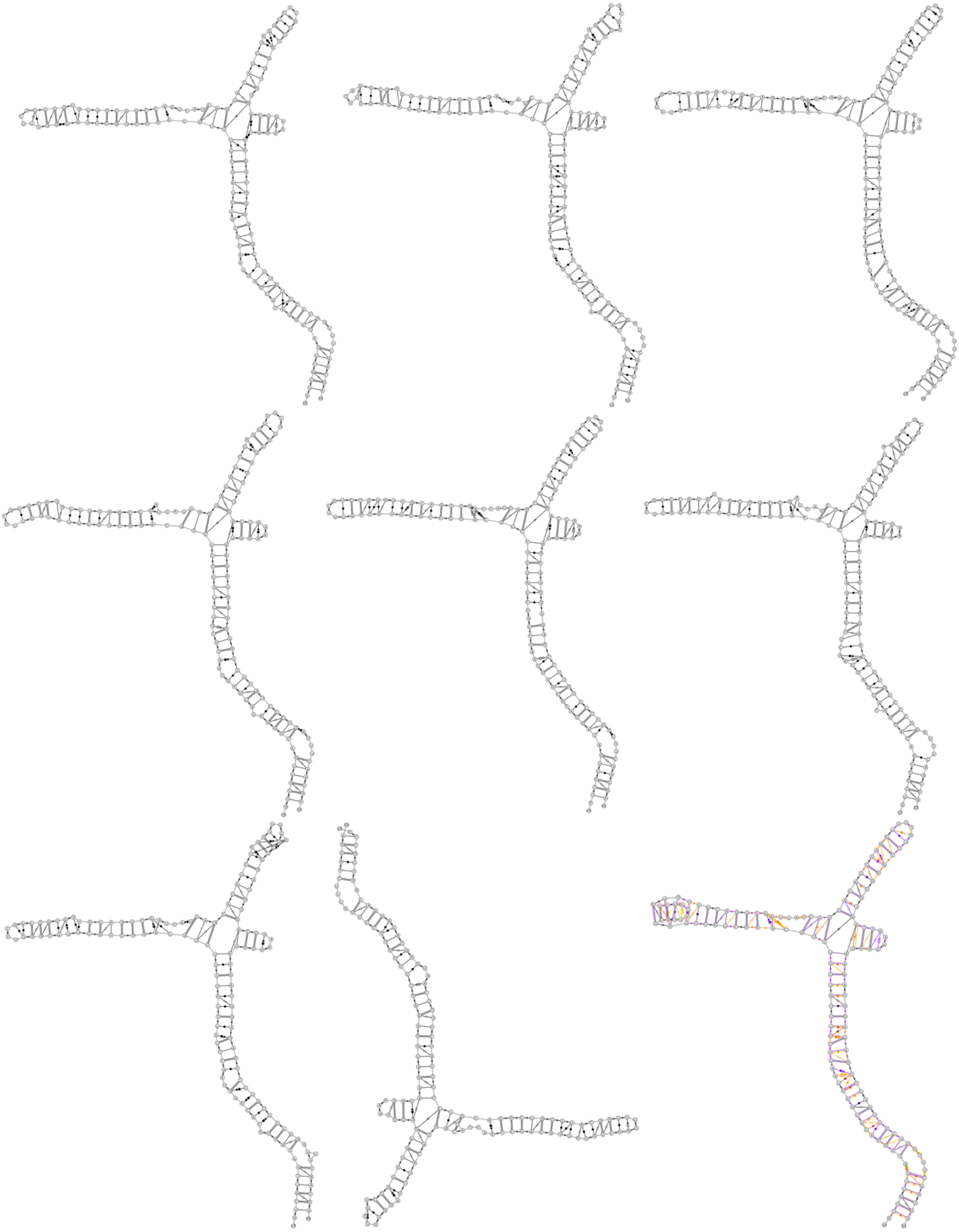

**Figure S23.** Barnaba representation of the states and of the full length simulation for P enantiomers with SL5

### f. SL3 RNA (no cylinders)

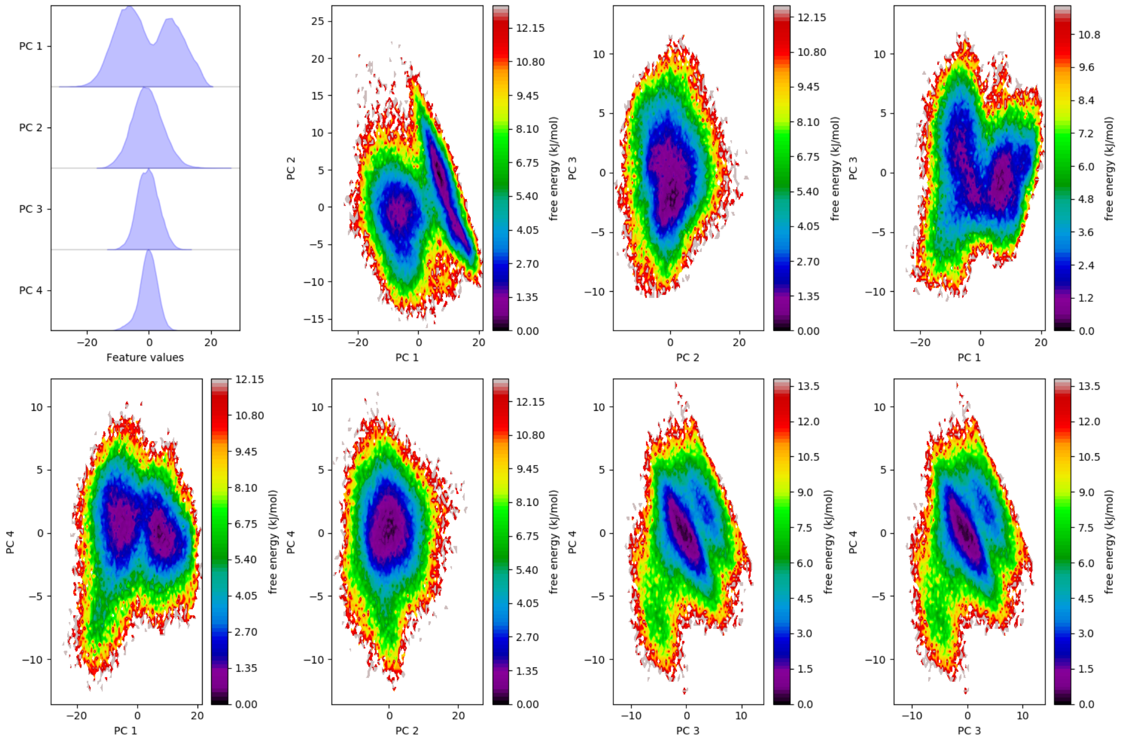

**Figure S24.** combined PCA of SL3 simulations (4μs)

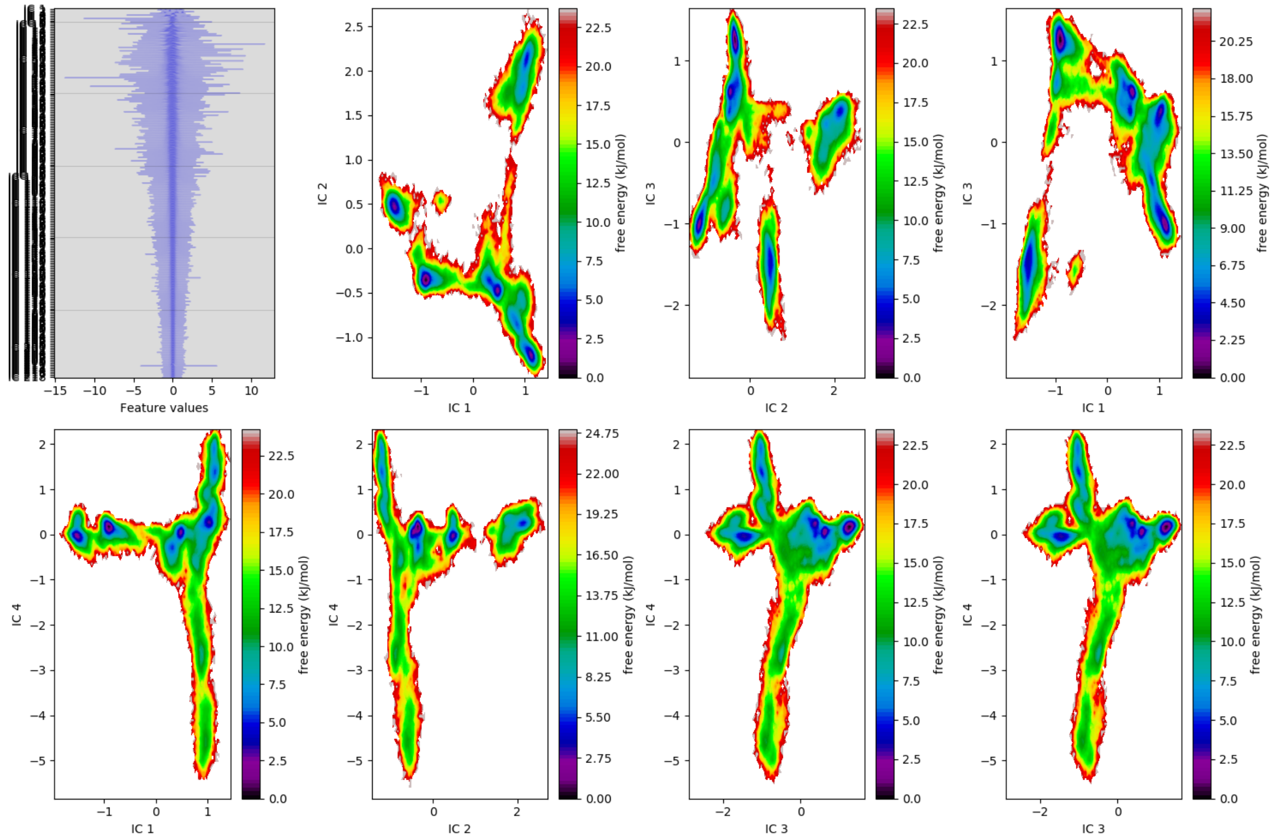

**Figure S25.** combined TICA of SL3 simulation

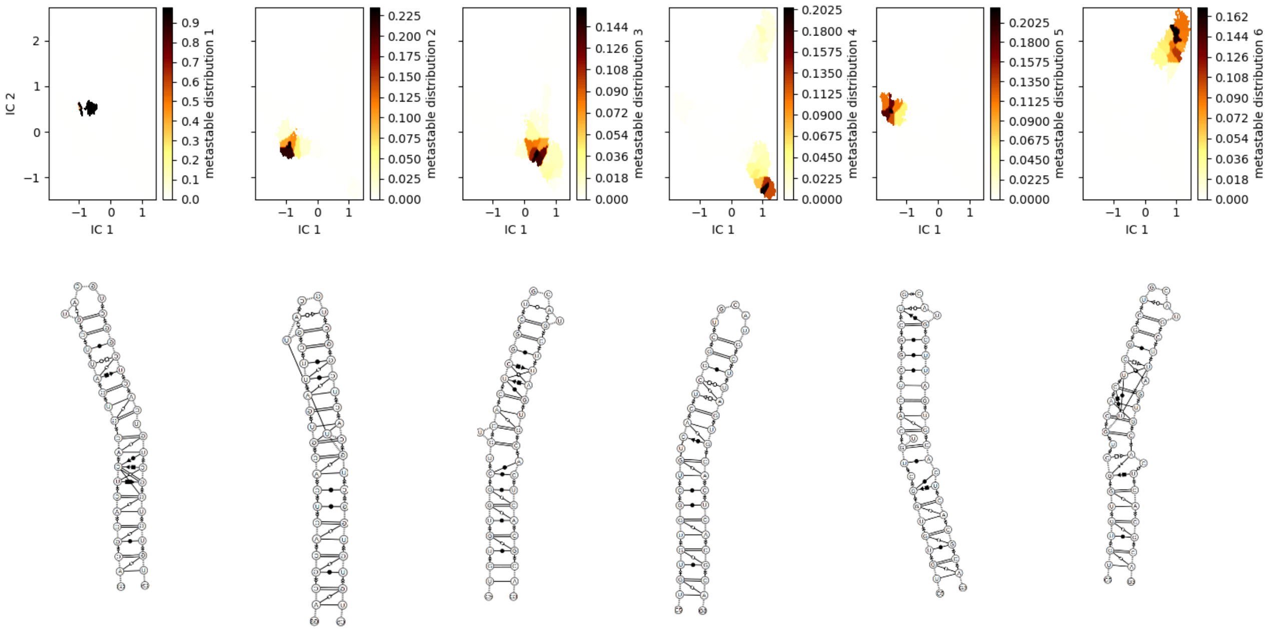


**Figure S26.** Metastable state separation of the TICA landscape, and corresponding SL3 structure
